# Unraveling MLL1-fusion Leukemia: Epigenetic Revelations from an iPS Cell Point Mutation

**DOI:** 10.1101/2022.12.16.520790

**Authors:** Laila Kobrossy, Weiyi Xu, Chunling Zhang, Wenyi Feng, Christopher E. Turner, Michael S. Cosgrove

## Abstract

Our understanding of acute leukemia pathology is heavily dependent on 11q23 chromosomal translocations involving the mixed lineage leukemia-1 (MLL1) gene, a key player in histone H3 lysine 4 (H3K4) methylation. These translocations result in distinct MLL1-fusion (MLL1F) proteins that are thought to drive leukemogenesis. However, the mechanism behind increased H3K4 trimethylation in MLL1F-leukemic stem cells (MLL1F-LSCs), following loss of catalytic SET domain of MLL1 (known for H3K4 mono- and dimethylation), remains unclear. In our investigation, we introduced a homozygous loss-of-function point mutation in MLL1 within human induced pluripotent stem cells. Remarkably, this mutation mimics the histone methylation, gene expression, and EMT phenotypes of MLL1F-LSCs- without the need for a translocation or functional wild-type MLL1. This observation underscores the essential role of MLL1’s enzymatic activity in restraining the cascade of epigenetic changes associated with the gene activating H3K4 trimethylation mark, which we show is catalyzed by mislocalized SETd1a H3K4 trimethyltransferase in the absence of MLL1’s enzymatic activity. Challenging existing models, our findings imply that MLL1F-induced leukemias arise from a dominant-negative impact on MLL1’s histone methyltransferase activity. We advocate for a therapeutic paradigm shift, targeting SETd1a for precision medicine. This work opens new avenues for addressing the complexities of MLL1-associated leukemias and improving targeted therapies.

**Summary:** Epigenomic and gene expression changes in iPS cells with a mutated MLL1 histone methyltransferase suggest an oncogenic mechanism for MLL1-fusion leukemias.

## Introduction

Understanding the formation of leukemic stem cells (LSCs) and the regulatory enzymes governing genome accessibility is crucial for addressing aberrant gene expression in leukemia. The mixed lineage leukemia family (MLL1-4 and SETd1a,b), pivotal in histone H3 lysine 4 methylation (H3K4me1,2,3), shapes chromatin architecture critical for the transcriptional programs that maintain cell identity ^1–5^.

Chromosomal translocations involving the MLL1 gene associate with poor-prognosis acute leukemias, fostering an aberrant transcriptional plasticity or permissiveness that results in a mixed lymphoid/myeloid cellular identity ^6–8^. These translocations in one allele of MLL1 produce MLL1- fusion proteins (MLL1F) that replace the catalytic SET domain with one of over 80 fusion partners that are thought to drive leukemogenesis ^9^. Despite the structural and functional diversity of MLL1F proteins, common among MLL1F-LSCs is a hyper-H3K4 trimethylation phenotype that results in a substantial upregulation of LSC maintenance genes, including HOXA9, MEIS1, MEF2C, and ZEB1^10–19^. This investigation sought to explain a paradox in the oncogenic mechanism of MLL1F leukemias: How removal of the catalytic SET domain from a complex that predominantly catalyzes H3K4 dimethylation results in upregulated H3K4 trimethylation and target gene expression in MLL1F-leukemic stem cells.

H3K4 methylation plays a critical role in gene expression for neurogenesis, hematopoiesis, and embryonic development ^1–5^. Different methylation states of H3K4—mono-, di-, or trimethylation—associate with distinct enzymatic complexes and functions ^20^. H3K4me3, found in active gene promoters ^21–23^, recruits nucleosome remodeling complexes and promotes RNA PolII promoter-proximal pause release ^24–29^. H3K4me2 is associated with poised transcription and, unexpectedly, with transcriptional repression ^24,30^. H3K4 monomethylation can activate or silence gene expression, depending on its location^31–35^.

Despite known hyper-H3K4 trimethylation in MLL1F-LSCs^16,19,36^, the role of MLL1 enzymatic alterations remains elusive. Prevailing theories suggest that MLL1F proteins interact with components of transcriptional elongation machinery that together activate the H3K4 trimethylation activity of wild-type MLL1 encoded from the unaffected allele ^37–40^. However, this model doesn’t account for various scenarios, including fusion proteins that don’t interact with the elongation machinery ^9^, partial tandem duplications of MLL1 associated with AMLs that silence the unaffected allele ^41^, or the loss of both MLL1 alleles in some patient samples ^42^. In contrast, conditional deletion of the MLL1 SET domain doesn’t alter H3K4 methylation levels in MLL1-AF9 cells in mice, suggesting dispensability for leukemogenesis^43^. However, the possibility that MLL1F proteins function in a dominant-negative manner by disrupting the enzymatic activity of the protein encoded from the unaffected allele^44^ remains unclear.

Current models also overlook detailed biochemical, structural, and evolutionary analyses showing product specificity differences among human MLL1 paralogs (see **Fig. *S1***). Isolated SET domains from MLL1 family members primarily catalyze slow H3K4 monomethylation, requiring interaction with the WRAD (<u>W</u>DR5, <u>R</u>bBP5, <u>A</u>sh2L and <u>D</u>PY30) sub-complex for recognition of the nucleosome substrate and for di- and/or trimethylation^45–48^ (**Fig. 1a**, and **Fig. *S1&S2***). Complexes assembled with MLL1 and MLL4 (also known as MLL2^49^) predominantly catalyze mono- and dimethylation, while MLL2 (also known as MLL4) and MLL3 focus on monomethylation ^48^. In contrast, SETd1a,b complexes catalyze mono-, di-, and trimethylation^48^ (**Fig. *S1***). The mechanism underlying the loss of MLL1’s catalytic domain leading to increased trimethylation in LSCs remains enigmatic.

**Fig. 1.**
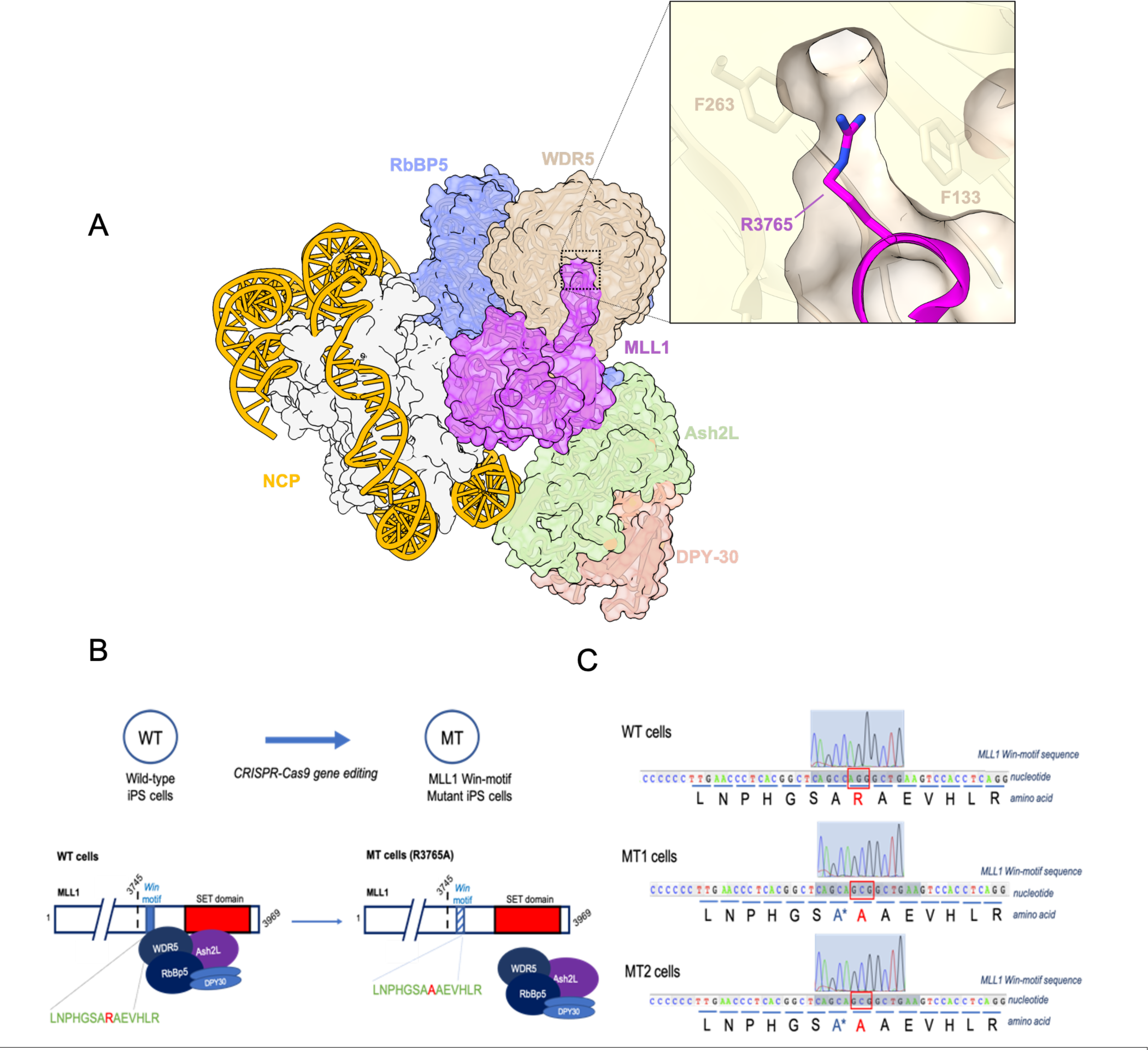
Generating a homozygous R3765A point mutation of a key arginine residue in the MLL1 Win-motif critical for MLL1 core complex assembly and enzymatic activity. a, Three-dimensional structure of the MLL1 core complex bound to a nucleosome core particle (NCP) with DNA in yellow and histones white (PDB code: 7UD5). Inset: cross-section of the X-ray structure of WDR5 (tan) bound to MLL1 Win motif peptide (magenta) (PDB code: 4ESG). The positions of the crucial Win motif arginine 3765 in MLL1 and WDR5 residues F133 and F263 are indicated. b, Upper: schematic of mutant (MT) iPS cell line generation from wild-type (WT) cells using the CRISPR-Cas9 system. Lower: loss-of-function of MLL1 Win-motif through arginine>alanine point mutation leads to complex disassembly and loss of H3K4 methylation activity (Supp. Fig. 1). c, Chromatograms of sequencing results confirming R3765A mutation in exon 32 of the KMT2A locus in MT cells. Two clones, MT1 and MT2, were generated. Genomic DNA from WT (upper), mutant clone MT1 (middle), and clone MT2 (lower) iPS cells was extracted and PCR used to amplify the MLL1 Win-motif region. * To prevent recutting the modified genome after repair, a silent mutation GCC>GCA was introduced in the single-stranded oligodeoxynucleotide donor site that was the DNA template for homology-directed repair.

Our investigation explored the consequences of a homozygous point mutation causing the loss of histone methyltransferase activity in MLL1 within human induced pluripotent stem (iPS) cells. Surprisingly, this genetic alteration led to a cascade of effects, unveiling an abnormal pattern of histone methylation, altered gene expression, and a cellular phenotype remarkably reminiscent of MLL1F leukemic stem cells. Intriguingly, these parallels emerged in the absence of an MLL1 translocation or a functional copy of wild-type MLL1, shedding light on the critical role of MLL1’s dimethylation activity in maintaining cell identity, with implications for our understanding of the mechanism of MLL1F leukemogenesis.

A pivotal revelation from our study is the profound impact of MLL1’s dimethylation activity in constraining the cascade of epigenetic changes linked with the activating H3K4 trimethylation mark. This process is essential for curbing aberrant epigenetic plasticity and safeguarding cellular identity. Our findings provide compelling evidence that the observed hypertrimethylation phenotype arises from the mislocalization of a different enzyme, the SETd1a H3K4 trimethyltransferase at crucial gene promoters maintaining LSCs.

This discovery challenges conventional perspectives and introduces a novel interpretation: MLL1F-induced leukemias likely originate from a dominant-negative impact on MLL1’s histone methyltransferase activity, like the consequences observed here with a homozygous loss-of-function mutation in the MLL1 gene within iPS cells. Consequently, our results suggest that targeted interventions focusing on MLL1 may inadvertently promote LSC formation, potentially challenging detection due to their slow growth characteristics. Instead, we advocate for a paradigm shift towards targeting SETd1a for inhibitor development as a more favorable approach to counteract abnormal H3K4 trimethylation in MLL1F leukemias. This strategic redirection in therapeutic targeting opens new avenues for precision medicine in addressing the intricacies of MLL1-associated leukemias.

## Results

### Generating a homozygous loss-of-function point mutation in MLL1 in human iPS cells

We previously demonstrated the dependence of the MLL1 core complex’s enzymatic activity on WDR5’s recognition of Arg3765, situated within the highly conserved WDR5 interaction (Win) motif of MLL1 (**Fig. 1a** (inset); and **Fig. *S2a***) ^50–53^. The X-ray structure reveals that the guanidinium moiety of R3765 inserts into the central tunnel of the seven-blade WD-repeat structure of WDR5, stabilized by cation-Pi interactions with WDR5 residues F133 and F263 (**Fig. 1a**, inset)^52^. Substitution of R3765 with alanine abolishes MLL1 core complex assembly *in vitro* and *in cellulo (***Fig. *S2b,c***), resulting in a nearly complete loss of H3K4 methylation activity with histone peptides or the more physiological nucleosome substrate ^50,54–56^ (**Fig. *S2b,d***).

Small molecules mimicking the Win motif, collectively referred to as Win motif inhibitors or *Wini*, bind WDR5 and hinder MLL1 core complex assembly and enzymatic activity in vitro ^50,54,57^. They selectively antagonize primary and immortalized MLL1F and CEBPA-mutated leukemia cell lines ^58,59^, as well as cell lines with gain-of-function P53 mutations that alter MLL family enzyme expression ^60^. Notably, this antagonism does not extend to other cell lines lacking MLL1 translocations, CEBPA mutations, or gain-of-function P53 mutations^58–60^.

However, the role of MLL1’s enzymatic activity in leukemia pathogenesis and as a potential target for inhibition remains controversial. While there is a suggestion that specific inhibition of the MLL1- WDR5 interaction is the mechanism underlying selective antagonism of MLL1F leukemias ^58^, this model doesn’t account for the conservation of the Win motif among all family members and the possibility that other family members may be targeted. Indeed, we have shown that MLL1 and SETd1a complexes, but not those of other family members, are equally sensitive to *Wini* molecules ^57^. More broadly, our understanding of the role of MLL1’s enzymatic activity in cellular identity and differentiation has been hindered by the common use of retroviral transduction of MLL1F proteins into aneuploid immortalized cell lines containing wild-type copies of MLL1^61^, along with the frequent use of methylation state-specific antibodies with significant cross-reactivity issues^62^.

To avoid these ambiguities, CRISPR-Cas9 was used to introduce the homozygous R3765A loss-of-function point mutation in the endogenous MLL1/KMT2A gene locus in karyotypically normal human ASE-9203 iPS cells. We targeted a 20 bp region in the MLL1/KMT2A gene to introduce an R3765>A3765 point mutation (AGG>GCG) in both alleles of the KMT2A exon 32 (**Fig. 1b,c**). Two single cell-derived clones, 14 (MT1) and 39 (MT2), with homozygous mutations were identified and confirmed by in-house DNA sequencing (**Fig. 1c**). Sequencing also confirmed the absence of unintended mutations in the *Win*-motifs of the other five MLL/SET1 family members and eliminated the possibility of off-target effects in the top 10 predicted CRISPR/Cas9 off-target loci (**Fig. *S3a,b***). Isogenic wild type cells (WT) were used as the control.

### MLL1 R3765A substitution increases H3K4 trimethylation at transcription start sites

To investigate the impact of the *Win*-motif mutation on global H3K4 methylation levels in MT cells, we used western blots with whole cell extracts from wild type (WT) and both MT clones. The loss-of-function mutation might have caused a global decrease in H3K4 methylation levels in MT cells ^15,63^. Instead, we observed differential effects on each methylation state. H3K4 monomethylation and dimethylation levels in both MT clones were similar to WT, as were global levels of total histone H3 (**Fig. 2a**). Unexpectedly, both MT1 and MT2 showed *increased* levels of trimethylated H3K4 compared to WT (**Fig. 2a**). Likewise, immunofluorescence showed similar increases in H3K4 trimethylation in both MT clones compared to WT (**Fig. 2b**), suggesting that loss of MLL1 enzymatic activity leads to increased H3K4 trimethylation throughout the genome. Since a similar H3K4 hypertrimethylation defect was noted in mouse MLL1-AF10 LSCs (with AF designating the fusion partner) ^19^ and in patient-derived MLL1- AF4 blast cells ^36^, these results suggested that the enzymatic activity of MLL1 may be required to limit the spread of the gene-activating H3K4 trimethyl mark.

**Fig. 2.**
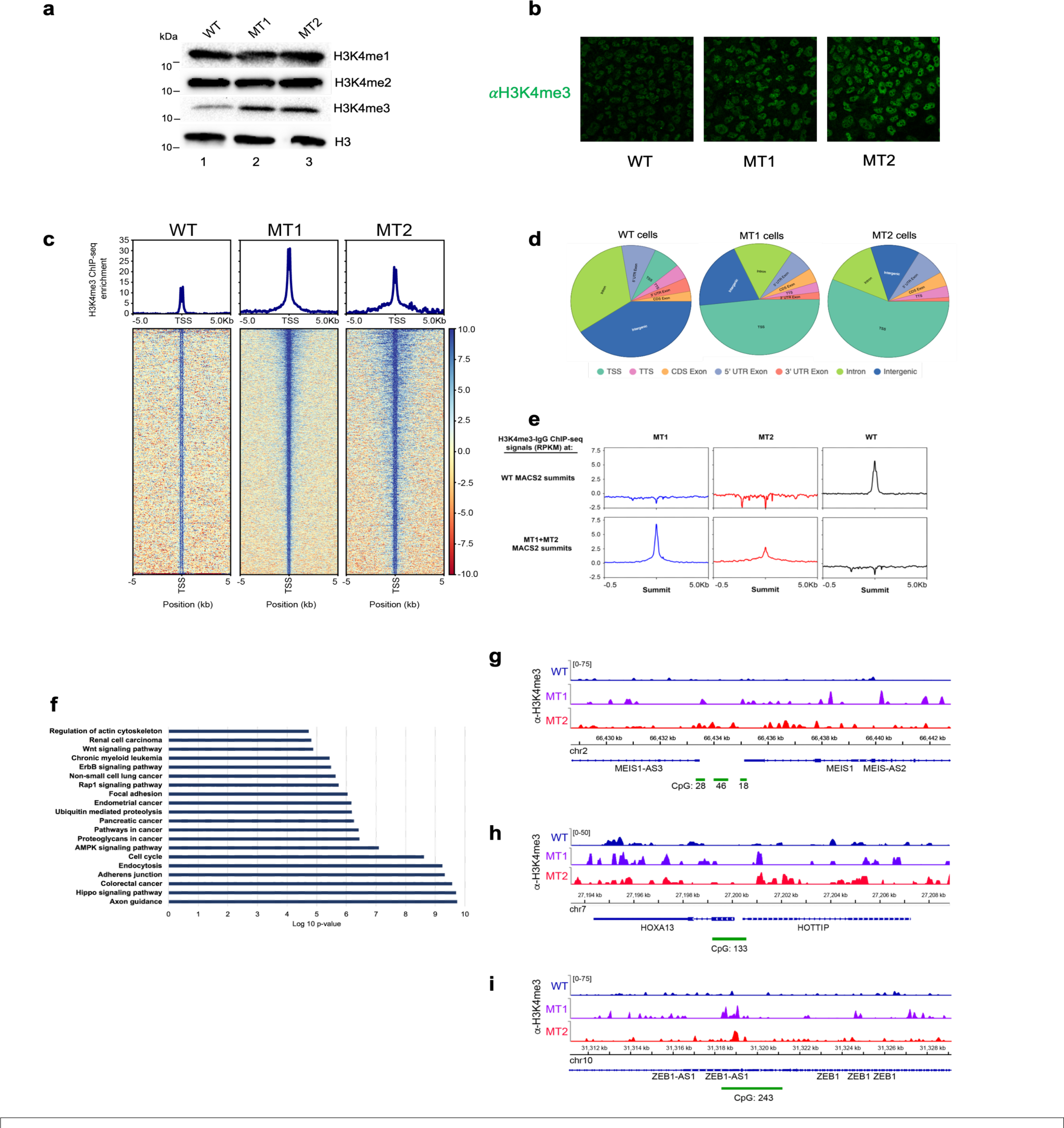
**Loss of MLL1 enzymatic activity results in a global increase and redistribution of H3K4 trimethylation. a**, Immunoblot analysis of H3K4me1-3 levels in WT and MT hIPS cells. Total H3 was used as the loading control. **b,** Immunofluorescence staining with anti-H3K4me3 antibody of WT and MT hIPS cells. **c,** ChIP-seq heat maps and profiles were generated from H3K4me3 – control IgG signals and plotted over transcription start sites (TSS +/- 5kb) for WT and MT hIPS cells. **d,** Pie charts for H3K4me3 enrichment vs. IgG in annotated genomic regions for WT and MT hIPS cells. **e,** Metagene plots for H3K4me3 ChIP-seq signal (-IgG) at WT MACS2 peak summits (top row) and at MT1 + MT2 MACS2 peak summits (bottom row). **f.** KEGG enrichment analysis of top canonical pathways represented in overlapping ChIP-Seq peaks from MT1 and MT2 clones normalized to WT. Log 10 of enrichment p-values (p< 0.05) for each pathway is represented on the x-axis. **g,h,i,** Integrated genomics viewer (IGV) browser snapshots comparing H3K4me3 – IgG ChIP-seq enrichment signals between WT and MT hIPS cells at the Meis1 promoter (g), HoxA13-HOTTIP promoter (h), and the Zeb1 promoter (i).

To further investigate this hypothesis, we performed chromatin immunoprecipitation followed by sequencing (ChIP-seq) to determine how loss of MLL1 enzymatic activity affects H3K4 trimethylation across the genome in iPS cells using a SNAP-ChIP validated anti-H3K4me3 polyclonal antibody (Epicypher, 13-0041)^62^. Similar to observations in MLL1-AF10 LSCs ^19^ and patient-derived MLL1-AF4 leukemia cells ^36^, we observed an increase in localization and spreading of H3K4 trimethylation peaks around transcriptional start sites in both MT1 and MT2 cells compared to WT (**Fig. 2c,d**). Peak annotation and gene enrichment analysis revealed significant overlap in the genomic regions of MT1 and MT2 cells that acquired novel H3K4 trimethylation peaks compared to WT (**Fig. *S3c***). Furthermore, principal component analysis (PCA) shows clustering of both mutants (**Fig. *S3d***), confirming consistent changes in the two biological replicates. These results suggest that there is a significant increase in the H3K4me3 at new genomic loci in the MT IPS cells. To further test this hypothesis, we plotted H3K4me3 ChIP-seq signals observed in each strain around the H3K4me3 peak summits observed in WT (**Fig. 2e**, top row), and around peak summits observed in both MT strains (**Fig. 2e**, bottom row). The results clearly show a redistribution of the H3K4me3 mark to new genomic loci in both MT strains compared to WT.

Kyoto Encyclopedia for Genes and Genomes (KEGG) enrichment analysis of the 5091 overlapping H3K4 trimethylation peaks in MT1 and MT2 cells revealed a significant enrichment for genomic regions encoding biomarkers of multiple cancer networks (see **Fig. S3e**), WNT signaling molecules, cell cycle machinery, cell adhesion, and the epithelial-mesenchymal transition (EMT) signaling pathway (**Fig. 2f**). Strikingly, increased H3K4 trimethylation occurred in the promoters of several of the same genes that are frequently upregulated in MLL1F leukemias including HoxA9-13, Meis1, and ZEB1 (**Fig. 2g-i**). Interestingly, increased trimethylation was also observed in the promoter of the lncRNA HOTTIP, which has been shown to coordinate activation of several HoxA genes from the 5’ tip of the HoxA cluster (**Fig. 2h**).

Taken together, these results suggest that loss of MLL1 enzymatic activity results in a genome wide increase in H3K4 trimethylation, especially in the promoters of LSC maintenance genes, and a potential shift to a more open, transcriptionally permissive chromatin state. Furthermore, that this phenotype occurs in a homozygous MLL1 loss-of-function strain demonstrates that the hyper-H3K4 trimethylation phenotype observed in MLL1F LSC’s can occur without a functional copy of wild type MLL1.

### MLL1 R3765A substitution increases expression of MLL1F LSC maintenance genes

To determine if the changes in global H3K4 trimethylation levels were associated with changes in gene expression, we used RNA-seq to compare transcriptional dynamics of MT and WT cells and found ∼2800 differentially expressed genes (**Fig. 3a**). Consistent with a more permissive chromatin environment, the majority of genes were upregulated in MT compared to WT cells (1914 genes, or 68%) (**Fig. 3a**). Gene Set Enrichment Analysis revealed that of the 25 highest scoring (most similar) gene sets, about 50% were associated with aggressive malignancies and multicancer invasiveness (**Fig. 3b**). The other half correlated with gene sets involved in chromatin remodeling (25%), extracellular matrix dynamics and EMT signaling (17%), and stem cell pluripotency (8%). The relevance of these signaling cascades in cancer was confirmed via ingenuity pathway analysis, as highlighted in the top 10 canonical pathways differentially regulated in MT vs. WT iPS cells (**Fig. 3c**).

**Fig. 3.**
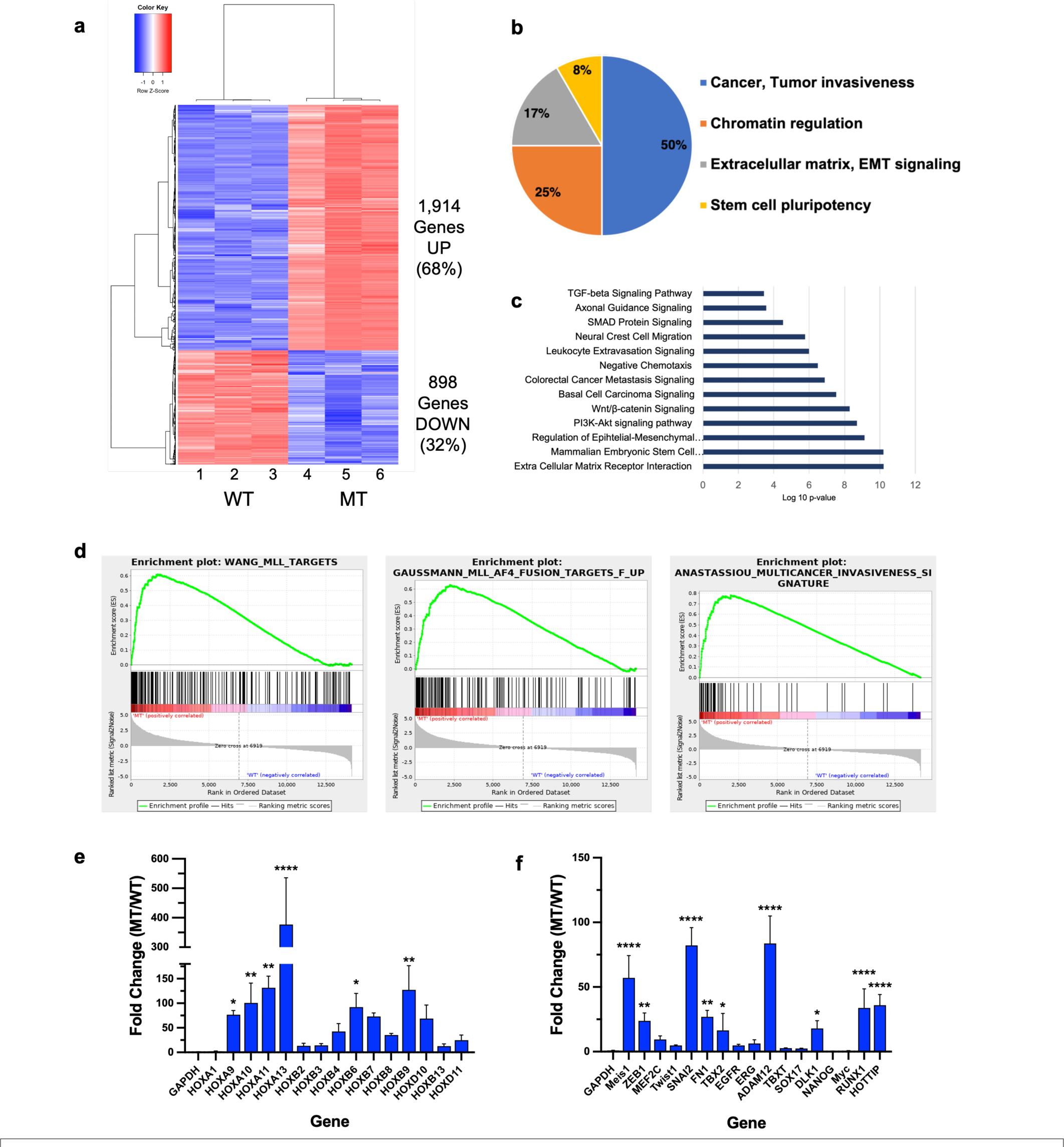
**Loss of MLL1 enzymatic activity increases expression of LSC maintenance genes. a**, Heatmap showing differentially expressed genes in MT vs. WT cells (clone MT2) in RNA-Seq. The heatmap was generated using normalized counts from three independent replicates of total RNA from WT (columns 1-3) and MT cells (columns 4-6). **b,** Pie chart showing the composition of the 25 highest scoring gene sets for differentially expressed genes in WT vs. MT hIPS cells. **c,** Gene ontology enrichment analysis showing top canonical pathways in differentially expressed genes in MT cells. X-axis, log10 p-values; y-axis, list of top overrepresented pathways based on all differentially expressed genes in MT vs. WT cells. **d,** Representative Gene Set Enrichment Analyses (GSEA) for differentially expressed genes in MT vs. WT hIPS cells. **e,f,** Mean fold change (MT/WT) in mRNA expression levels for select LSC maintenance genes; including: **e,** HOXA and HOXB cluster genes, and **f,** other oncogenes commonly associated with MLL1F leukemias. Error bars represent std. dev. from three independent experiments. The relative change in GAPDH expression was included as a control on each plot. (****p<0.0001; ***p<0.001, **p<0.01, *p<0.05)

Further evaluation of the cancer-specific gene sets revealed significant enrichment of MLL1 target genes ^15^, MLL1F-induced gene expression patterns ^64^, and cancer invasiveness transcriptional signatures ^65^ (**Fig. 3d**). Consistent with the ChIP-seq results, we observed significant upregulation of several leukemia- and stem cell-associated genes including HOX genes and cancer associated microRNAs, many of which are key players in the development of MLL1F leukemias ^66–73^ (**Fig. 3e,f**). For example, expression of Hox genes A9, A10, A11 and A13 increased from ∼77- to 377-fold (**Fig. 3e**), in a manner associated with their chromosomal distance from the HOTTIP lncRNA gene at the 5’ end of the HoxA with previous studies showing HOTTIP functions in locus control through a chromatin looping mechanism that spatially coordinates expression at the 5’ end of the HoxA cluster ^74^. Indeed, increased HOTTIP expression has been shown to increase HoxA gene expression and AML-like disease in mice and is associated with poor prognosis MLL1F-AML in humans^75^. Furthermore, a ∼20-100-fold increase in expression of several genes associated with MLL1F leukemias was observed, including Meis1, RUNX1, ZEB1, HOXB6, and ADAM12 ^13,76–78^(**Fig. 2e,f**).

These observations were not due to differences in MLL1 mRNA or protein levels (**Fig. S3a,b**), or differences in WDR5 and RbBP5 protein levels in MT cells (**Fig. S3b**). These results point to a shift to a more permissive epigenetic state that supports aberrant transcriptional activation of key regulatory genes in MT cells. Furthermore, the occurrence of this aberrant gene expression signature in cells without an MLL1F protein suggests that it is the loss of MLL1 enzymatic activity that drives overexpression of the core genes associated with MLL1F leukemias.

### MLL1 R3765A promotes a “mixed lineage” differentiation phenotype in iPS cells

Aberrant expression of multiple lineage markers in the same cell during hematopoietic differentiation exemplifies the “mixed lineage” phenotype in MLL1F leukemias ^79,80^. To determine if loss of enzymatic activity of MLL1 in the absence of a fusion protein contributes this phenotypic plasticity, we investigated if MT cells could differentiate into the three germ layers (endoderm, mesoderm, and ectoderm) compared to WT. We used commercially available differentiation media that are used to assess stem cell pluripotency and qPCR to compare expression of lineage markers Sox17 (Endoderm), TbxT (Mesoderm) and Dlk1 (Ectoderm) before and after differentiation (**Fig. 4a**). WT cells differentiated into each of the three germ layers, as demonstrated by the significantly increased expression of lineage-specific makers in differentiated WT cells compared to undifferentiated controls (**Fig. 4b-d**). MT cells also differentiated into each of the three germ layers, with a significantly higher potency for endoderm and mesoderm layers, as shown by the relative fold-increase in gene expression in MT cells compared to WT cells. However, differentiating MT cells consistently displayed increased transcriptional or epigenetic plasticity, resulting in a “*mixed-lineage*” phenotype in endoderm and mesoderm germ layers (**Fig. 4b,c**), demonstrated by the misexpression of two or more lineage-specific markers in cells treated with single-lineage-specific growth media. In contrast, ectoderm differentiation was unaffected by the mutation (Fig. 4d). Since it has been shown that WDR5 expression decreases upon ESC differentiation ^81^, differences in the initial differentiation state of the cells cannot explain the observed differentiation phenotypes since similar levels of the WDR5 protein were observed in WT and both MT cells (**Fig. S4b** and **Fig. 6a** below).

**Fig. 4.**
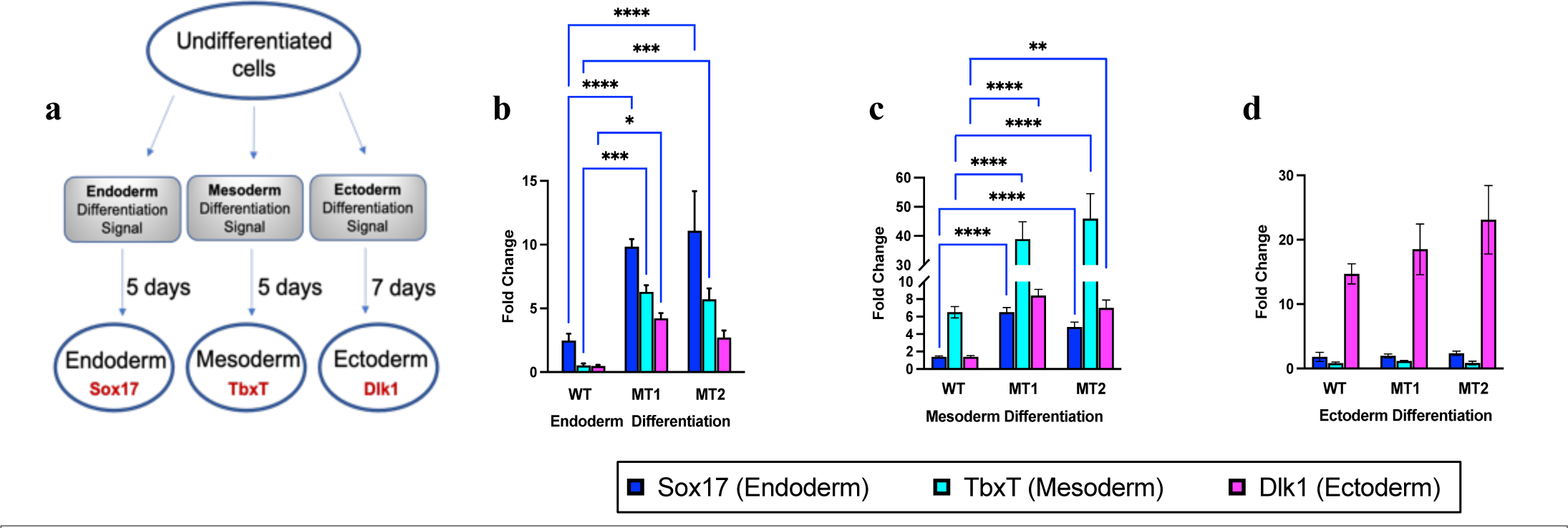
**R3765A substitution in MLL1 induces a “mixed lineage” phenotype in iPS cells. a**, Undifferentiated WT and MT (MT1 and MT2 clones) iPS cells were treated with STEMdiff endoderm, mesoderm, or ectoderm differentiation media for 5-7 days, followed by RNA extraction and RT-qPCR analysis for expression of endoderm (Sox17), mesoderm (TbxT), and ectoderm (Dlk1) lineage-specific biomarkers. **b,c,d**, WT and MT gene expression changes upon differentiation in (**b**) edoderm, (**c**) mesoderm, and (**d**) ectoderm lineages after incubation of cells with the indicated STEMdif media. Mean fold-increase in biomarker gene expression relative to undifferentiated controls was quantified by RT-qPCR using cDNA collected at experimental endpoints (Day 5 for endoderm and mesoderm, Day 7 for ectoderm), and after normalization to GAPDH levels. Error bars represent the standard deviation from three technical replicates for each experiment (****p<0.0001; ***p<0.001, **p<0.01, *p<0.05).

These results suggest a tissue specific role for the enzymatic activity of the MLL1 core complex in conveying a precise tissue-specific patterning across endoderm and mesoderm germ layers, possibly by limiting epigenetic plasticity at key MLL1 target genes.

### MLL1 R3765A substitution promotes EMT and increases invasiveness

The activation of the epithelial-mesenchymal transition (EMT) pathway is a well-established hallmark of cancer invasion and metastasis, particularly in the context of cancer stem cell development and maintenance ^82–84^. Specific transcription factors, including Zeb1, Twist1, and Slug, mediate the EMT pathway, leading to the repression of epithelial-specific adhesion molecules such as E-cadherin ^85,86^. Concurrently, there is transcriptional activation of mesenchymal structural components (N- cadherin, vimentin) and extracellular matrix molecules like fibronectin ^87–90^. Notably, transcripts of several mesenchymal signature biomarkers were significantly elevated in MT cells (**Fig. 5a**) and considering their association with EMT and poor survival in MLL1-AF9 AML patients ^18^, we aimed to investigate whether the loss of MLL1 activity was linked to EMT-associated cancer stem cell formation and increased invasiveness.

**Fig. 5.**
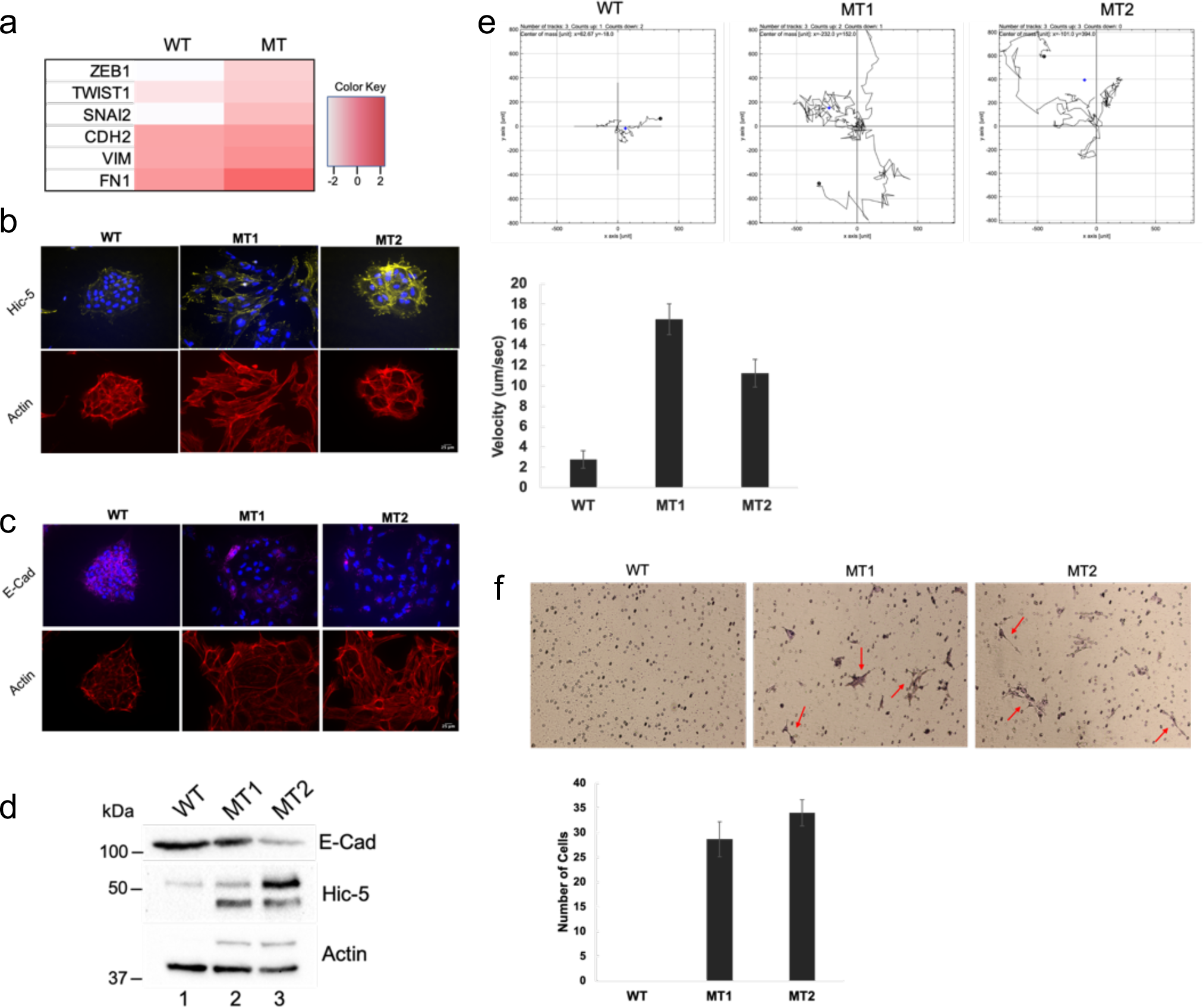
R3765A substitution in MLL1 induces EMT and invasiveness. a, Heatmap of mesenchymal signature biomarkers differentially expressed in RNA-Seq of mutant (MT) vs. wild-type (WT) cells. Significantly altered transcripts were selected based on adjusted p-value and FDR ≤ 0.05. Color key shows differential expression based on a log2 scale. **b,** Immunofluorescence of WT and MT1 and MT2 cells showing differential staining for Hic-5 (yellow). DAPI (blue), nuclear regions; phalloidin (red) actin filaments. **c,** Immunofluorescence of WT and MT1 and MT2 cells showing differential staining for E-cadherin (E-Cad, magenta). DAPI (blue), nuclear; phalloidin (red), actin filaments. **d,** Immunoblots of whole cell extracts from WT (lane 1) and MT1, lane 2, and MT2, lane 3 cells using antibodies specific for E-cadherin (E-Cad) and Hic-5. Beta-actin was the loading control. **e,** Upper: Rose diagrams showing cell motility assays in WT and MT1 and MT2 cells by time-lapse microscopy imaging. Lower: Bar chart of cell migration velocity. Error bars are standard deviation with p-value 0.05. **f,** Upper: Photographs of representative Matrigel invasion assays, performed as two biological replicates. Red arrows, Giemsa-stained invading MT cells. Lower: Cells counted in three separate vision fields for each biological replicate, with average number of invading cells/view for each cell type. Error bars are 1 standard deviation with p-value 0.05.

**Fig. 6.**
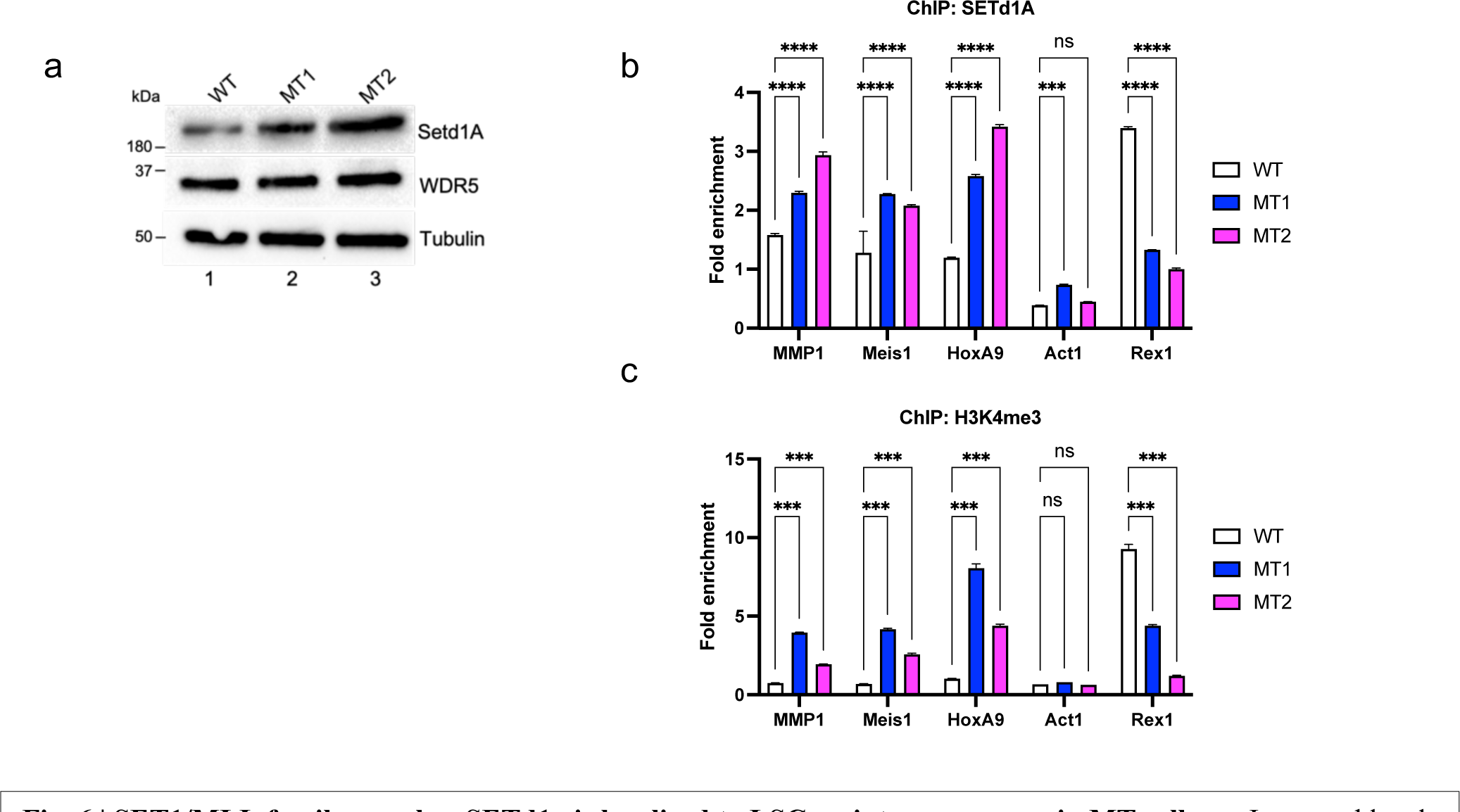
SET1/MLL family member SETd1a is localized to LSC maintenance genes in MT cells. a, Increased levels of SETd1A in mutant (MT) cells. Immunoblot analysis was performed on whole cell extracts from wild-type (WT) (lane1) and MT (MT1, lane2; MT2, lane 3) cells using antibodies specific for SETd1a and WDR5. Tubulin was the loading control. **b,** ChiP-qPCR results for SETd1a localization to promoters of upregulated genes (MMP1, HOXA9, MEIS1), a downregulated gene (Rex1p) and a control gene with unchanged expression in WT vs. MT cells. Fold-enrichment was relative to an IgG ChIP control for each sample, and all values were normalized to the GAPDH housekeeping gene. **c,** as in (b) except ChIP used an H3K4 trimethylation antibody. Error bars represent the standard deviation from three technical replicates for each experiment (****p<0.0001; ***p<0.001, **p<0.01, *p<0.05).

To explore the newly acquired mesenchymal and oncogenic characteristics of MT cells, we initially compared MT and WT cells for the protein levels and localization of E-cadherin and Hic-5, signature biomarkers for epithelial and mesenchymal cells, respectively ^85,91,92^. Immunohistochemistry revealed increased staining at focal adhesions of the promigratory, adhesion adapter protein Hic-5 (**Fig. 5b**), and lower levels of epithelial cell-specific E-cadherin at the cell periphery in both MT clones ompared to WT (**Fig. 5c**). Phalloidin-conjugate staining exposed actin stress fibers in both MT clones (**Fig. 5b**), suggesting a potential shift in cytoskeletal and cell migration dynamics. These findings were corroborated by immunoblots, indicating a reduction in E-cadherin protein in both MT clones, accompanied by a corresponding increase in Hic-5 (**Fig. 5d**).

To determine if cell movement dynamics in MT cells were typical of cells that have undergone an EMT, we conducted time-lapse imaging of WT and MT cells and characterized cell motility parameters including migration velocity. Motility assays revealed a significant increase in migration of both MT clones compared to WT iPS cells (**Fig. 5e**), further suggesting increased invasive potential with the loss of MLL1 activity.

We next investigated if MT cells acquired invasive characteristics, as suggested by the RNA-seq and ChIP-seq pathway enrichment. In transwell invasion assays, multiple MT cells invaded extracellular matrix-coated filters (**Fig. 5f**), suggesting substantial invasive potential. In comparison, no control WT iPS cells invaded, as expected from nondifferentiated iPS cells with more epithelial cellular characteristics, including robust cell-cell adherens junctions ^86^ (**Fig. 5b**). These results were consistent with the hypothesis that MLL1’s histone methyltransferase activity is important for restricting the EMT phenotype, and loss of activity promotes cancer stem cell formation and invasiveness.

### MLL1 R3765A promotes redistribution of SETd1a trimethyltransferase to LSC gene promoters

MLL1F leukemia cells show increased H3K4 trimethylation that is thought to result from the MLL1F protein binding to MLL1 target genes and activating wild type MLL1 expressed by the unmutated allele of these heterozygous cells ^37^. However, the similar ectopic H3K4 trimethylation phenotype in MT cells with a homozygous loss-of-function point mutation indicated involvement of a different enzyme. Humans have 6 MLL family members: MLL1-4, SETd1a and SETd1b. Biochemical reconstitution revealed that SETd1a and SETd1b are efficient trimethyltransferases responsible for the bulk of H3K4 trimethylation in cells while MLL1-4 are relatively poor trimethyltransferases ^48,93,94^. Furthermore, small hairpin RNA knockdown of SETd1a but not SETd1b transcripts induces differentiation and apoptosis in murine and human MLL1F-leukemia cells ^95^. We therefore hypothesized that increased H3K4 trimethylation is due to relocalization of SETd1a to MLL1 target gene promoters.

Consistent with this hypothesis, both MT clones had increased SETd1a protein relative to WT (more noticeable in MT2) (**Fig. 6a**). In contrast, WDR5 levels were unchanged in both MT clones (**Fig. 6a**). ChIP-qPCR with SETd1a and H3K4 trimethylation antibodies determined if their localization was altered at promoters of MLL1 target genes in MT cells. SETd1a (**Fig. 6b**) and H3K4 trimethylation (**Fig. 6c**) levels increased at promoters of genes for MMP1, Meis1 and HoxA9, which RNA-seq showed were all highly upregulated in MT cells. In contrast, the REX1 promoter showed decreased SETd1a localization and the ACT promoter was relatively unchanged, consistent with, respectively, downregulated and unchanged mRNA expression by RNA-seq. These results suggested a key role for SETd1a relocalization and its associated H3K4me3 activity in creating a more permissive chromatin environment that promotes the mixed lineage cancer stem cell phenotype after loss of MLL1 enzymatic activity.

## Discussion

Despite thirty years of research into the molecular basis of the “mixed lineage” phenotype observed in cells with MLL1 translocations^79,80^, prognostic challenges persist in leukemias with MLL1 mutations^96^. The intricate interplay involving MLL1’s histone methyltransferase activity complicates our understanding of its oncogenic mechanism. The existence of a noncatalytic fusion partner replacing the SET domain in one MLL1 allele in MLL1F cells suggests a potential haploinsufficiency mechanism.

However, the paradoxical increase in H3K4 trimethylation, driving the expression of crucial LSC maintenance genes, adds layers of complexity.

Our study, introducing a homozygous loss-of-function point mutation in MLL1, phenocopies the MLL1F LSC phenotype. We observe a widespread increase in H3K4 trimethylation at LSC target genes, accompanied by their significant upregulation in gene expression compared to wild-type cells. The “mixed lineage” phenotype during differentiation, resembling MLL1F cells, and altered expression of epithelial- mesenchymal transition genes underscore the similarity to LSCs. Importantly, these phenotypes manifest in the absence of an MLL1-fusion protein or a functional copy of wild-type MLL1, suggesting that it is the loss of MLL1’s enzymatic activity that is the crucial driver of the key epigenomic changes underlying cellular transformation in MLL1F leukemias (**Fig. 7**).

**Fig. 7.**
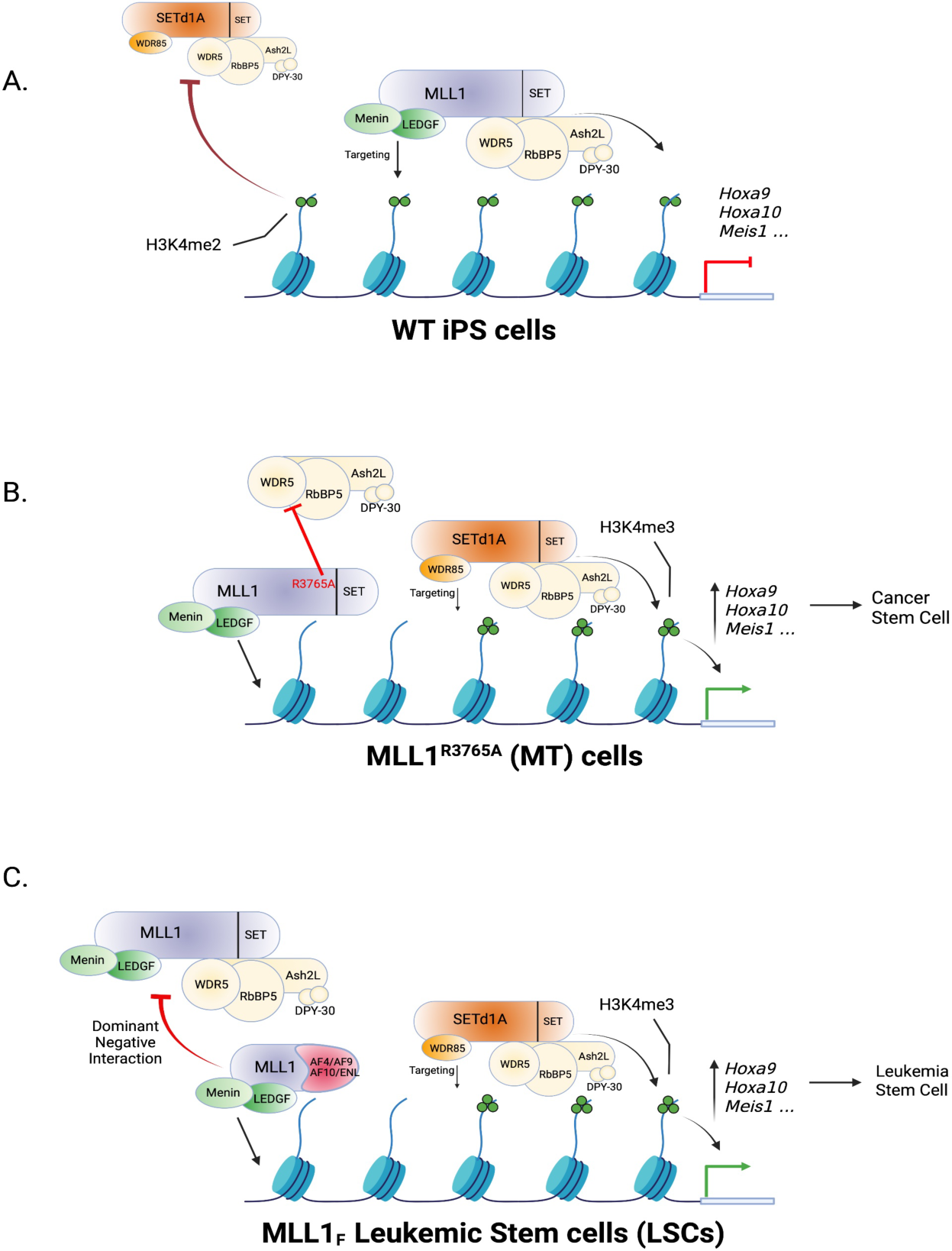
Model for the role of MLL1 histone methyltransferase activity in normal and leukemic stem cells. a, Wild type (WT) MLL1-catalyzed H3K4 dimethylation (H3K4me2) limits the spread of the gene-activating H3K4 trimethylation (H3K4me3) mark catalyzed by SETd1a in WT iPS cells. **b**, Homozygous loss-of-function point mutation of MLL1 prevents deposition of the H3K4me2 mark at MLL1 target genes. This allows spreading of the SETd1a-catalyzed H3K4me3 mark and activation of LCS maintenance genes Hoxa9, HoxA10, Meis1, etc. **c,** Model for MLL1F-induced leukemogenesis. In this model, the MLL1F protein acts in a dominant negative manner byinteraction with or displacement of WT MLL1 and prevents H3K4me2 methylation at MLL1 target genes. As in the homozygous loss-of-function point mutant cells (b), lack of MLL1 catalytic activity allows spreading of the SETd1a-catalyzed H3K4me3 mark to MLL1 target gene promoters resulting in expression of LSC maintenance genes. Created with BioRender.com.

These findings are consistent with the previously proposed dominant negative mechanism in MLL1F leukemias, where MLL1F proteins, lacking the catalytic SET domain, inhibit the enzymatic activity of wild-type MLL1 encoded from the unaffected allele^44,97^. Taking a step further, we posit that the loss of MLL1 enzymatic activity is a unifying feature of leukemias with and without MLL1 translocations, including AML resulting from a partial tandem duplication in the MLL1 gene, where it has been shown that expression of the wild type allele is epigenetically silenced in patient MLL1-PTD leukemic blasts ^98^, and in patient samples in which both MLL1 alleles have been deleted^42^.

This conclusion contrasts with other studies suggesting that wild type MLL1 is required for interaction of the MLL1F protein with LSC target genes^37^ or that MLL1’s enzymatic activity is dispensable for leukemogenesis^43^. These investigations focused on the impact of knockdown or conditional deletion of wild-type MLL1 in the presence of an MLL1 fusion protein, overlooking the potential dominant negative role of the fusion protein on MLL1’s enzymatic activity. Little or no change in H3K4 methylation and HOX gene expression was observed in these studies, consistent with a dominant negative mechanism in which the enzymatic activity of MLL1 is already inhibited by the fusion protein. Furthermore, recent evidence shows that the MLL1-AF6 fusion protein does not require wild type MLL1 for persistent localization to chromatin, demonstrating that wild-type MLL1 is not necessary for binding of the oncoprotein^99^. A dominant negative mechanism also explains why an MLL1 N-terminus fused to the transcriptionally inert LacZ protein produces acute myeloid leukemia when knocked into the MLL1 locus in mice ^100^ and is consistent with studies showing specific sequences from the MLL1-N terminus and/or the fusion protein are required for cellular transformation ^101–105^.

Moreover, a loss of function mechanism may explain the occurrence of cancer-associated nonsense and missense mutations in the MLL1 gene in cancer databases. Supporting a potential tumor suppressor function, our recent findings demonstrate that up to 85% of the 29 cancer-associated missense mutations localized in or around the MLL1 SET domain are defective for histone H3K4 mono- and/or dimethylation^106^. This further supports the notion that the enzymatic activity of MLL1 plays a crucial role in preventing aberrant cellular transformations, aligning with its function as a tumor suppressor. This also suggests that loss of MLL1 enzymatic activity may play a wider role in solid tumor development than previously appreciated.

Therapeutically, strategies targeting the MLL1F-menin interaction or stabilizing wild-type MLL1 levels gain significance in the context of a dominant negative mechanism. MLL1F proteins require interaction with the menin tumor suppressor for binding MLL1 target genes^107^. Disruption of this interaction, as proposed by various inhibitors^108^, is consistent with counteracting the inhibitory effects of MLL1F proteins acting in a dominant negative manner. Stabilizing wild-type MLL1 levels, in approaches inhibiting its proteasomal degradation, aims to restore the balance disrupted by MLL1F proteins^109^, providing further evidence for a dominant negative interplay.

Crucially, our findings shed light on the relationship between MLL1 and SETd1a, the H3K4 trimethyltransferase complex. Biochemical reconstitution experiments demonstrate that human SETd1a/b complexes efficiently catalyze H3K4 trimethylation^48^ and are crucial for the bulk of this modification in mammalian cells^93,94^. We show that the loss of MLL1 enzymatic activity results in the redistribution of SETd1a and the H3K4 trimethyl mark to the promoters of select LSC maintenance genes. This implies that MLL1’s mono- and dimethyltransferase activity may limit the spread of the gene-activating H3K4 trimethylation mark catalyzed by SETd1a. Intriguingly, H3K4 dimethylation is associated with recruitment of the NCoR/SMRT histone deacetylase corepressor complex in humans and in yeast^24,30^. Loss of NCoR/SMRT localization, combined with relocalization of SET1a, may underlie increased histone acetylation and H3K4 trimethylation in the absence of MLL1’s H3K4 dimethylation activity in MLL1F leukemias. This model is also consistent with the demonstration that knockdown of SETd1a, but not other MLL family members, antagonizes growth of murine MLL1-AF9 and human MOLM-13 leukemic cells ^95^.

In summary, our investigation reveals a sophisticated cancer model where the loss of MLL1 enzymatic activity, facilitated by a dominant negative mechanism, remodels the chromatin landscape. This alteration results in increased epigenetic plasticity, enabling the selection of more differentiated progeny cells that elude normal growth controls, exhibit enhanced invasiveness, and acquire drug resistance more readily^110^. The proposed interplay between MLL1 and SETd1a sheds light on how MLL1 influences H3K4 trimethylation dynamics. From a therapeutic standpoint, targeting the H3K4 trimethylation activity of the SETd1a complex emerges as a promising strategy for MLL1F-leukemias and other cancers with MLL1 mutations. This underscores the critical importance of understanding the intricate mechanisms governing epigenetic regulation in leukemia progression.

## Acknowledgements and Funding

The authors would like to thank Ashley Canning and Michael Connelly for helpful comments on the manuscript and Anne Smardon for helpful discussions. We thank Steve Hanes and Leszek Kotula for qPCR instrument access, and Ryan Palumbo and Baylee Porter for help with qPCR experiments. We thank Karen Gentile at SUNYMAC core facility for her help with ChIP-Seq and RNA-seq data collection, and Frank Middleton for help with data deposition. We also thank Elizabeth Duncan, Saima Rahman, and the journal club trainees at the University of Kentucky Markley Cancer Center for helpful comments on our BioRxiv manuscript.

This work was supported by NIH R01 CA140522 to M.S.C., NIH R35 GM131709 to C.E.T., NIH R21 MH130812 to W.F., and a Richard & Jean Clark Pediatric Research Funds Award to M.S.C. and C.E.T.

## Data Availability

ChIP-seq and RNA-seq datasets have been deposited in the NCBI GEO database under accession numbers GSE213239 and GSE213237, respectively.

## Declaration of AI or AI-assisted technologies in the writing process

During the preparation of this work, the authors used ChatGPT Version 3.5 to edit for brevity and clarity. After using this tool, the authors reviewed and edited the content as needed and take full responsibility for the content of the publication.

## Materials and Methods

### Generating the MLL1 Win-motif homozygous mutation in human iPS cells

ASE9203 human iPS cells (WT cells) were from Applied Stem Cell (ASC), and CRISPR-Cas9-mediated gene editing was by ASC using their proprietary CRISPR-Cas9 protocol. Puromycin-resistance screening was performed on single cell-derived colonies and two clones (MT1 and MT2) homozygous for the MLL1 Win motif R3765A were identified and sequenced.

### Cell Culture

Both WT and MT iPS cells were maintained in mTeSR^TM^1 medium with 1x Supplement media (Stem Cell Technologies, Catalog #85850) and supplemented with 10 μM Y27632 (Stemgent) only upon thawing and passaging. Cells were plated and maintained on Matrigel hESC-qualified Matrix (Corning) according to the Stem Cell Technologies “Maintenance of Human Pluripotent Stem Cells in mTeSR^TM^1” manual. Passaging was with ReLeSR passaging reagent (Stem Cell Technologies). Due to their highly adhesive characteristics, MT cells were periodically gently scraped when passaged to release them from Matrigel-coated wells, and passaged at higher density (1:3 dilution per passage) relative to WT cells (1:10 dilution per passage). Cells were maintained for a maximum of three passages due to growth constraints of MT cells following the third passage.

### Tri-lineage Differentiation

WT and MT cells were propagated separately in lineage-specific differentiation media for endoderm, mesoderm or ectoderm using the STEMdiff Trilineage Differentiation Kit Protocol (Stem Cell Technologies). After differentiation, total RNA was extracted from duplicate cultures, and qRT-PCR was performed to evaluate expression of lineage-specific signature markers for endoderm (Sox17), mesoderm (TbxT), or ectoderm (DLK1). Data were normalized to the housekeeping gene (GAPDH), and the ddCt algorithm was used for differential expression analysis and comparison between cell types.

### RNA Extraction and Reverse Transcription (RT)-qPCR

WT and MT cells were plated in 6-well Matrigel-coated plates (Corning) until 80-90% confluent. RNA was extracted using Trizol reagent (Invitrogen) according to manufacturer’s instructions. Reverse transcription used SuperScript IV Reverse Transcriptase (Invitrogen) using 1 ug total RNA per reaction and 5% of the synthesized cDNA was used in a 20 ul PCR reaction. Real-time qPCR ysed iQ SYBR Green Supermix (Biorad) in a Realplex 4 qPCR 96-Well Real Time Cycler (Eppendorf). Ct values were internally normalized to GAPDH for each sample, and the ddCt algorithm was used for differential expression analysis and comparison between cell type. Primer sequences are in the “Antibodies and Primers” section.

### RNA Sequencing

Whole transcriptome profiling was performed on technical triplicates of WT and MT cells (Clone MT2) at the SUNY Molecular Analysis Core (SUNYMAC) facility at Upstate Medical University. RNA was isolated using Trizol reagent. RNA quality and quantity were assessed with RNA 6000 Nano Kits on an Agilent Bioanalyzer 2100. Sequencing libraries were prepared with the TruSeq Stranded Total RNA Library Prep Kit RiboZero Gold (Illumina), using 1 ug total RNA as input. Library size was assessed with DNA 1000 Kits on an Agilent Bioanalyzer 2100, and libraries were quantified with Qubit dsDNA HS Assay Kits (Invitrogen). Libraries were pooled and sequenced on the NextSeq 500 instrument (Illumina), with paired end 2 x 75 bp reads using a High Output 150 cycle reagent kit.

An average of 54 M paired-end reads per sample was generated from sequencing. Sequencing quality was accessed by fastQC v0.11.8, low-quality bases/reads and adaptors were removed from reads by trimmomatic v0.38. Trimmed reads were mapped to the Gencode GRCh38 release 29 Human reference genome using STAR aligner v2.7.0d [3]. Reads mapped to genes were summarized by the featureCounts program in subread v1.6.3. Genes were filtered by CPM ≥1 in at least 2 samples, and data were normalized to effective library size using edgeR v3.22.5. Differential gene expression analyses were performed using edgeR programs. RNA-seq data have be deposited into the Gene expression Omnibus (GEO) database.

Genes with false discovery rate (FDR) ≤0.05 and fold-change ≥ 2 were considered significantly differentially expressed genes (DEGs) between MT and WT cells. DEGs were investigated for pathway enrichment by Ingenuity Pathway Analysis (IPA, Qiagen) and functional clustering by Database for Annotation, Visualization and Integrated Discovery (DAVID) v6.7. GSEA was performed on MT and WT transcriptome expression data using MSigDB database v6.2.

### Chromatin Immunoprecipitation

ChIP experiments were performed in duplicate for WT and MT cells (clones MT1 and MT2) using the High Sensitivity ChIP Kit (Abcam). 5x10^5^ cells were used as input material for each immunoprecipitation. Following crosslinking and cell lysis, DNA shearing was performed using Diagenode Bioruptor® Sonicator at high power output, with 5 pulses of 30 seconds, followed by 30 seconds rest between each pulse. For H3K4me3, 1 μg of Rabbit anti-H3K4me3 ChIP grade antibody (Epicypher, 13-0041), which has been SNAP-ChIP validated for H3K4me3 specificity ^62^, was used for the immunoprecipitation. Rabbit IgG (Thermo Fisher Scientific, 31235) (1 μg) was used as a negative control. Immunoprecipitated DNA was column purified and resuspended in 20μl DNA elution buffer. The purified DNA was either used for sequencing (ChIP-seq) or qPCR as indicated.

### Library Generation, Sequencing and Analysis for ChIP-Seq

ChIP-seq was performed on technical triplicates of WT and MT cells (Clones MT1 and MT2) at the SUNY Molecular Analysis Core (SUNYMAC) facility at Upstate Medical University. Due to low input material and therefore low yield following ChIP, all purified ChIP DNA was used as input for NEBNext Ultra II Library Prep Kits for Illumina. Library size was assessed with a DNA 1000 Kit on an Agilent Bioanalyzer 2100, and libraries were quantified with Qubit dsDNA HS Assay Kits (Invitrogen). Libraries were pooled and sequenced on a NextSeq 500 instrument (Illumina), with paired end 2 x 75 bp reads using High Output 150 cycle reagent kits. Processing and alignment of ChIP-seq data used Partek Flow analysis software. Reads were aligned using the BWA algorithm. ChIP peaks were called using MACS2 callpeak function with two-sample analysis between IP and IgG control (p value < 1e-5). Annotated peaks were used to generate TSS metaplots and for gene enrichment analysis. ChIP-seq metaplots were constructed using deepTools functions bamCompare and plotHeatmap^111^. Those MACS2 peaks from MT1 and MT2 cell lines were pooled. ChIP signals over the pooled MT MACS2 peaks or the WT MACS2 peaks were analyzed by the deepTools computeMatrix function, then visualized by the plotProfile function. Peak comparison analysis pipelines were generated with sample attributes specific for treatment (immunoprecipitated vs. IgG), genotype (WT vs. MT) and pair (MT1 and MT2 vs. WT). Annotated peaks were used to generate TSS metaplots and for gene enrichment analysis. Gene and pathway enrichment analyses used the KEGG Mapper online analysis tool, and Venn Diagrams were creating using the Bioinformatics & Evolutionary Genomics online tool (http://bioinformatics.psb.ugent.be/webtools/Venn/).

### ChIP-qPCR

Following ChIP, qPCR was performed using iQ SYBR Green Supermix (Biorad) in either a CFX Opus 384 Real-time PCR system (Biorad) or a realplex 4 qPCR 96-Well Real Time Cycler (Eppendorf) with 5% of the immunoprecipitated DNA used in a 20 ul PCR reaction. For both RT and ChIP, Ct values were internally normalized to GAPDH for each sample. For ChIP, fold-enrichment calculations for Setd1A and H3K4 trimethylation ChIP were normalized relative to IgG ChIP controls for each cell type. Each qPCR reaction was performed with technical triplicates. Primers set sequences are in the “Antibodies and Primers” section.

### Immunoblot Analysis

Whole cell extracts were prepared by lysing cells in RIPA buffer (50 mM Tris HCl pH 7.4/150 mM NaCl/0.5% sodium deoxycholate/0.1% sodium dodecyl sulfate/1% NP-40) containing cOmplete protease inhibitor cocktail (Roche), followed by quantification using Bradford assays. Approximately 20 ug protein was analyzed by 4%–15% gradient TGX Precast polyacrylamide gel (Bio-Rad), transferred onto a PVDF membrane (Biorad Transblot Turbo system) and probed for specific proteins (see “Antibodies and Primer Sequences” section). PVDF membranes were blocked for 1 hour with a 5% nonfat milk solution and incubated with primary antibody at 4°C overnight. Blots were washed three times with 0.1% TBS/Tween20 and incubated with HRP-conjugated secondary antibody for 1 hour at room temperature. Blots were washed three times and visualized by chemiluminescence (Clarity Western ECL Substrate, Bio-Rad) on a Bio-Rad ChemiDoc MP Imager using the chemiluminescence setting.

### Immunofluorescence

Cells on Matrigel-coated glass coverslips were fixed with 4% paraformaldehyde in PBS for 15 min, permeabilized with 1% Triton X-100 in phosphate buffered saline (PBS) for 15 min, quenched with 0.1 M glycine for 15 min and blocked with 3% bovine serum albumin (BSA) for 1 hour at room temperature. Coverslips were incubated with indicated primary antibodies diluted in 3% BSA for 2 hours at 37°C, followed by 1 hour incubation in fluorescent-tagged secondary antibodies as indicated. F-actin was stained with rhodamine-phalloidin (for cytoskeleton) and nuclei were stained with DAPI (Sigma-Aldrich). Cells were imaged using a Zeiss Axioskop2 plus microscope, fitted with a Q imagin ExiBlue charge-coupled device camera using an Apochromat 20Å∼ objective.

### Motility Assay

For live cell analyses, WT and MT iPS cells (clones MT1 and MT2) were plated in duplicate in 6-well tissue culture plates, coated with Matrigel and tracked using a Nikon TE2000 inverted microscope equipped with a temperature/CO2 regulated environmental chamber. Images were acquired at 10 min intervals for 14 hours. The manual tracking plugin of ImageJ was used to track the cell centroid over the duration of the movie. The Chemotaxis and Migration plugins of ImageJ were used to calculate velocity. Motility assays were performed on three independent biological replicates.

### Invasion Assay

Invasion assays were performed using Corning BioCoat 24-well Invasion Chambers (Corning). WT and MT cells were plated in duplicate in regular growth media (mTESR1 supplemented with 1x Supplement) at 5 x 10^4^cells/ml. Chemoattractant (mTESR1 supplemented with 5x Supplement media) was added to Falcon TC Companion plates, and chambers transferred to wells containing the chemoattractant. Cells were incubated overnight for 22 hours in a humidified tissue culture incubator at 37°C and 5% CO2 atmosphere. After incubation, non-invading cells were scrubbed from the upper surface of the chamber membranes using a cotton-tipped swab. Cells on the surface were washed twice with PBS, fixed with 100% methanol for 20 minutes at room temperature, washed twice with PBS, and stained with Giemsa (Sigma) for 20 minutes at room temperature. After final washes, cells were visualized under a light microscope and photographed with a Nikon microscope camera (FE 3291306). The total number of migrating cells was counted in three random vision fields per sample.

