## Supplementary information for "Unraveling MLL1-fusion Leukemia: Epigenetic Revelations from an iPS Cell Point Mutation"

Laila Kobrossy[a], Weiyi Xu[b,c], Chunling Zhang[d], Wenyi Feng[a], Christopher E. Turner[b], and Michael S. Cosgrove[a,*]

[a] Department of Biochemistry and Molecular Biology, SUNY Upstate Medical University, Syracuse, New York 13210, United States
[b] Department of Cell and Developmental Biology, SUNY Upstate Medical University, Syracuse, New York 13210, United States
[c] Current Address: Cell Biology Department, Harvard Medical School, Boston, MA
[d] Department of Neuroscience and Physiology, SUNY Upstate Medical University, Syracuse, New York 13210, United States

**Key Words:** Histone, methylation, leukemia, 11q23, translocation, epigenetic plasticity, H3K4 methylation, trimethylation, MLL, MLL1, KMT2A, SETd1A, KMT2F, EMT, WDR5, Win motif, cancer stem cell, leukemia stem cell, induced pluripotent stem cells, CRISPR-Cas9, ChIP-Seq, RNA-Seq, HOXA9, Meis1, HOTTIP, lncRNA, precision medicine

*To whom correspondence should be addressed:

Michael S. Cosgrove, Ph.D.
SUNY Upstate Medical University
Department of Biochemistry and Molecular Biology
750 East Adams Street
Syracuse, New York 13210


Supp. Fig. 1 and 2 are re-used from previous publications as indicated and are collected here to aid the reader in understanding the impact of the MLL1 R3765A substitution on the assembly and enzymatic activity of the MLL1 core complex *in vitro* and *in cellulo*.

#### Supp. Fig. 1 | The MLL1 core complex predominately catalyzes H3K4 mono- and dimethylation.

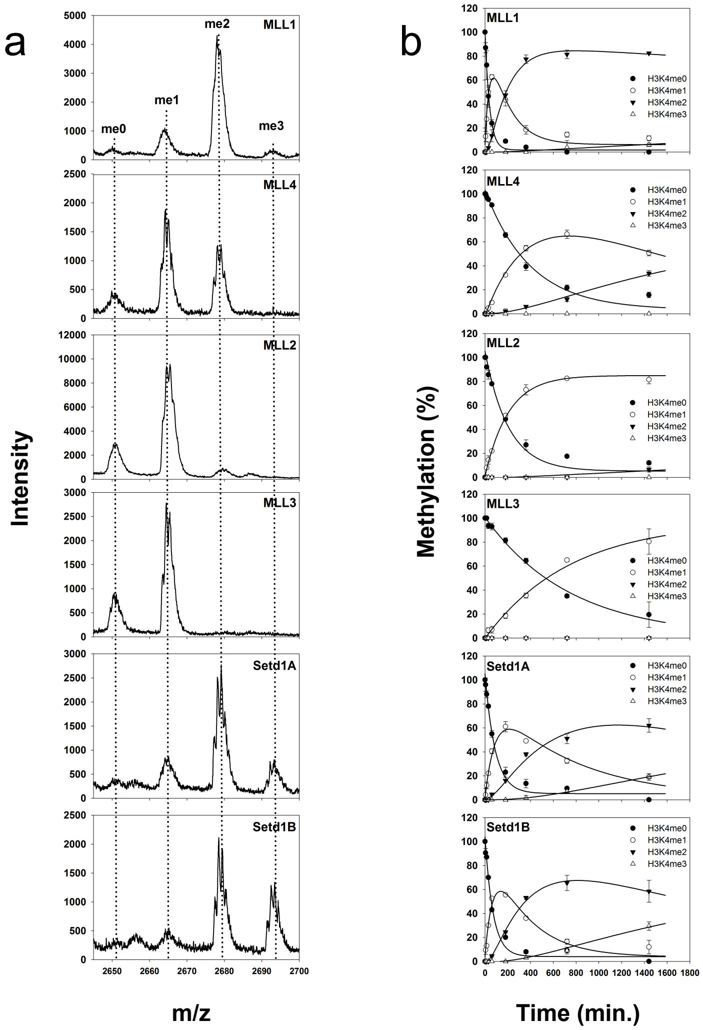


c

| Complex | Rate Constants ($hr^{-1}$)[a] | | |
|---|---|---|---|
| | $k_1$ | $k_2$ | $k_3$ |
| **MLL1 WRAD** | 1.50 ± 0.10 | 0.36 ± 0.04 | N/A[b] |
| **MLL4 WRAD** | 0.15 ± 0.009 | 0.04 ± 0.005 | N/A |
| **MLL2 WRAD** | 0.29 ± 0.02 | N/A | N/A |
| **MLL3 WRAD** | 0.08 ± 0.004 | N/A | N/A |
| **Setd1A WRAD** | 0.67 ± 0.05 | 0.08 ± 0.01 | 0.04 ± 0.01 |
| **Setd1B WRAD** | 0.85 ± 0.08 | 0.19 ± 0.03 | 0.03 ± 0.01 |

[a] Rate constants derived from global fitting of the data as described in methods ± standard error of the fit.
[b] N/A= non applicable, activities were either not present or too low to obtain reliable estimates of rate constants from global fitting.

**Fig. S1 | MLL/SET1 family core complex single turnover kinetics. a)** Quantitative MALDI-TOF mass spectrometry showing methylation of an unmodified histone H3 peptide (aa. 1-20) after 24 hours for each human MLL/SET1 family core complex. **b)** reaction progress curves globally fitted to irreversible consecutive reaction models using DynaFit. Each timepoint is the mean percentage of total integrated area for each species in MALDI-TOF reactions. Error bars represent +/- S.D. from duplicate measurements. **c)** summary of rate constants derived from global fitting of reaction progress curves. This figure is re-used from Fig. 3 of reference [5].

#### Supp. Fig. 1 | Cont.

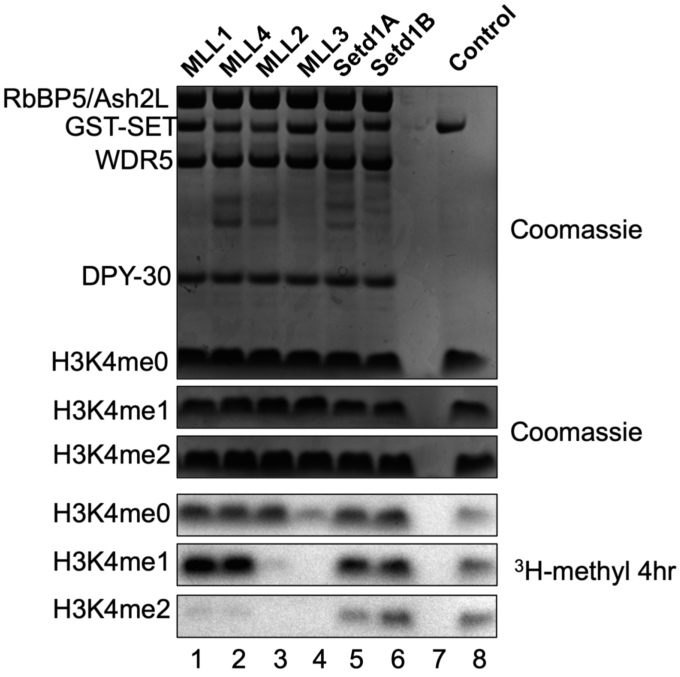


**Fig. S1 | Biochemically reconstituted human MLL/SET1 family core complexes have different product specificities.** Each MLL/SET1 family SET domain in a fusion with GST was incubated with WRAD subunits, $^{3}H$-methyl SAM, and histone H3 peptide substrates that were unmodified (H3K4me0), previously monomethylated at H3K4 (H3K4me1) or previously dimethylated at H3K4 (H3K4me2). Upper panels show Coomassie-Blue stained SDS-PAGE gels, lower panels show $^{3}H$-methyl incorporation after 4 hours as shown by fluorography. The control lane shows the activity of the isolated MLL1 SET domain on 100 µM unmodified H3 peptide, which was included on each gel. This figure is re-used from Fig. 2c of reference [5].

### Supp. Fig. 2 | The R3765A substitution in MLL1 disrupts the assembly and enzymatic activity of the MLL1 core complex.

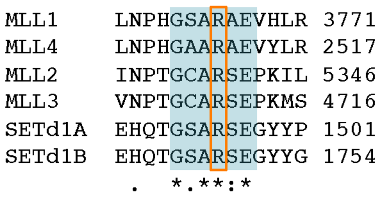


**Fig. S2a |** Clustal Omega multiple sequence alignment of human MLL1 family Win motif sequences (shaded in blue) with the critical arginine residue highlighted in the red box. This figure is re-used from Fig. 2A of reference [1].

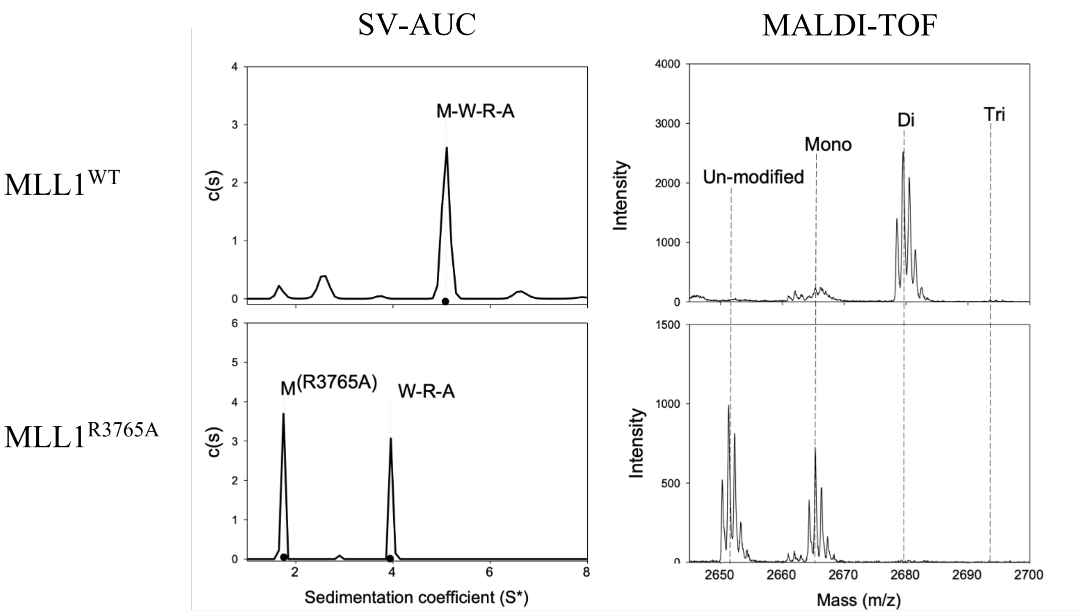


**Fig. S2b | The MLL1 R3765A substitution disrupts co-sedimentation of MLL1 with the WRA subcomplex and inhibits multiple methylation by the MLL1 core complex.** (Left column) Sedimentation coefficient distribution (c(s)) of the MLL1 core complex assembled with wild type MLL1 (residues 3745-3969) top panel, or with the R3765A MLL1 variant (bottom panel). SV-AUC, sedimentation velocity analytical ultracentrifugation; M, MLL1; W, WDR5; R, RbBP5; and A, Ash2L. (Right) MALDI-TOF mass spectra of quenched enzymatic reaction mixtures after 24 hours. The peaks show the relative amount of unmodified histone H3 peptide substrate (residues 1-20) that has been converted to H3K4me1,2,3 (mono-, di-, tri-) methylated species as indicated. This figure is re-used and adapted from Fig. 7 (panels A,B,G,H) of reference [3].

#### Supp. Fig. 2 | Cont.

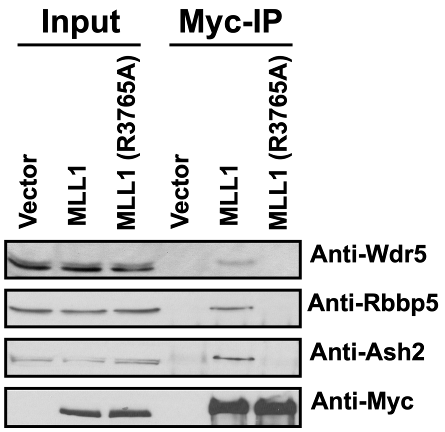


**Fig. S2c | The MLL1 R3765A substitution disrupts assembly of the MLL1 core complex in mammalian cells.** pCMV-Myc vectors encoding the 180 kDa C-terminal fragment of wild type or R3765A MLL1 proteins were transfected into HEK293 cells. Antibodies against c-Myc were used to immunoprecipitate each MLL1 variant. Pull downs were western blotted with anti-WDR5, anti-RbBP5 or anti-Ash2L antibodies. This figure is re-used from Fig. 2 of reference [2].

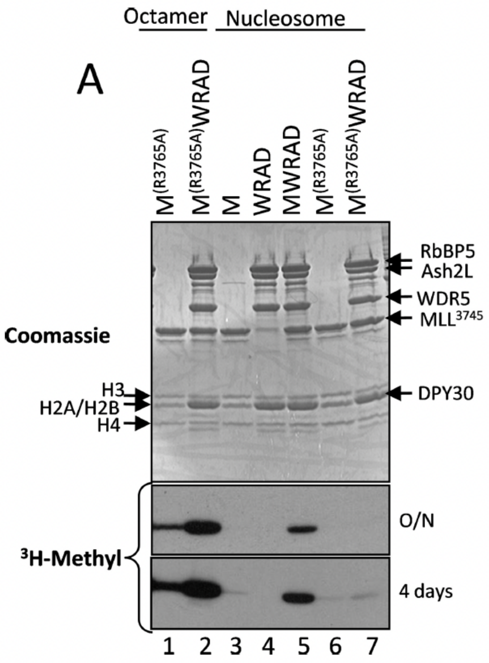


**Fig. S2d | The MLL1 R3765A substitution abolishes methylation of nucleosome histone H3 by the MLL1 core complex.** Wild type (lane 5) or R3765A (lane 7) MLL1 variants were assembled with WRAD and incubated with $^{3}$H-AdoMet and a reconstituted nucleosomal substrate. Quenched reactions were separated by SDS-PAGE and stained with Coomassie (upper panel). $^{3}$H-methyl signal was visualized by fluorography (lower panels). This figure is re-used from Fig 6A of reference [4].

#### Supp. Fig. 3 |

| MLL Family Win-motif Sequence Confirmation in WT and MT cells | | |
|---|---|---|
| Human MLL Family Member Gene (Protein) | DNA Coding Sequence | Amino Acid Sequence |
| KMT2D (MLL2) | atcaaccccactggctgtgccc**cga**tcagagcctaaaatcctc | GCARSE |
| KMT2C (MLL3) | cgttaaccccacaggttgtgccc**cgt**tctgaacctaaaatgagt | GCARSE |
| KMT2B (MLL4) | ctgaatccccatggggctgct**cgg**gcagaggtctatctccgg | GAARAE |
| KMT2F (Setd1A) | gagcaccagacaggctcagccc**cgc**agcgaaggctactacccc | GSARSE |
| KMT2G (Setd1B) | gagcacgtgacgggctgtgccc**cgc**agtgagggcttctacacc | GCARSE |

**Fig. S3a |** Integrity of all five Win-motif sequences for the other human MLL family members was confirmed in both WT and MT cells by Sanger sequencing. Win-motif coding sequence is in blue (key arginine coding sequences in bold).

| Off Target Locus | Sequence | PAM | Similarit y | Mismatch | Gene | Locus |
|---|---|---|---|---|---|---|
| 1 | GAGCCCTCAGGGCTCAGCCA | CAG | 6 | 3,10 | No | chr4@ 182905668-182905691 |
| 2 | GGACCCTGAGGGCTCAGCCA | AGG | 2 | 2,8,10 | No | chr15@ 74395949-74395972 |
| 3 | GGACCCTGAGGGCTCAGCCA | GGG | 2 | 2,8,10 | No | chr11@ 61518499-61518522 |
| 4 | GTCCCCTCACCGCTCAGCCA | GGG | 2 | 2,3,11 | No | chr16@ 689862-689885 |
| 5 | GACCACTCACGGCTCAGCCT | GGG | 1 | 3,5,20 | No | chr11@ 46384434-46384457 |
| 6 | GACCCCTCAGGGATCAGCCA | AGG | 1 | 3,10,13 | No | chr9@ 108810230-108810253 |
| 7 | GATCCATCACAGCTCAGCCA | CAG | 1 | 3,6,11 | No | chr4@ 8432046-8432069 |
| 8 | GAACCCACAAAGCTCAGCCA | GGG | 1 | 7,10,11 | No | chr6@ 52013917-52013940 |
| 9 | GAGCCCTCACAGCTCAGCCG | GGG | 1 | 3,11,20 | No | chr18@ 79803717-79803740 |
| 10 | GAGCCCTCACCCCTCAGCCA | AGG | 1 | 3,11,12 | No | chrY@ 17303630-17303653 |

**Fig. S3b |** Top 10 predicted CRISPR-Cas9 off-target loci identified by Applied Stem Cells. Sanger sequencing confirmed the integrity of all 10 loci in MT cells.

**Supp. Fig. 3 | Cont.**

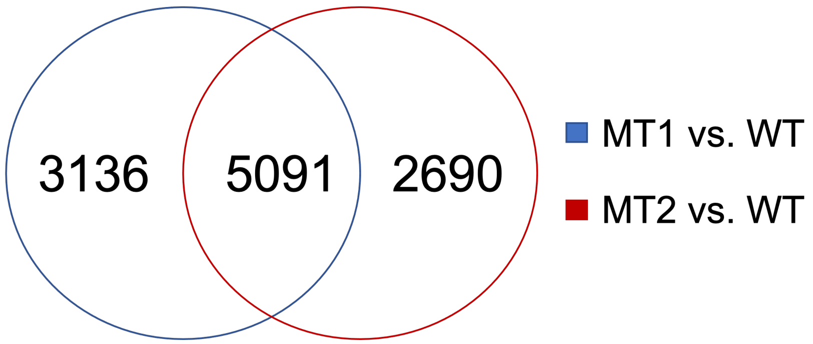


**Fig. S3c |** Venn diagram showing the overlap of differential ChIP-Seq H3K4me3 peaks detected in MT1 and MT2 clones, analyzed relative to WT cells.

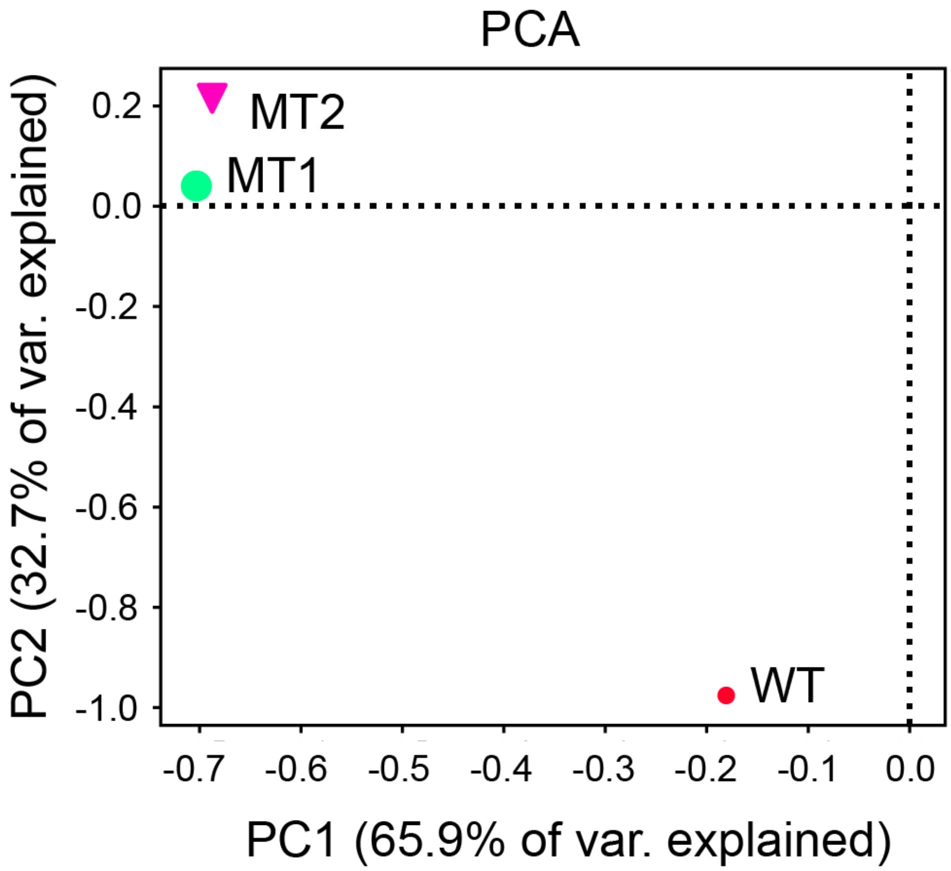


**Fig. S3d | Principal component analysis (PCA) of ChIP-seq results from Wild type and both MT IPS cell clones.** The grouping of both MT clones relative to WT confirms consistent changes in H3K4me3 ChIP-seq signals in the two MT clones.

**Supp. Fig. 3 | Cont.**

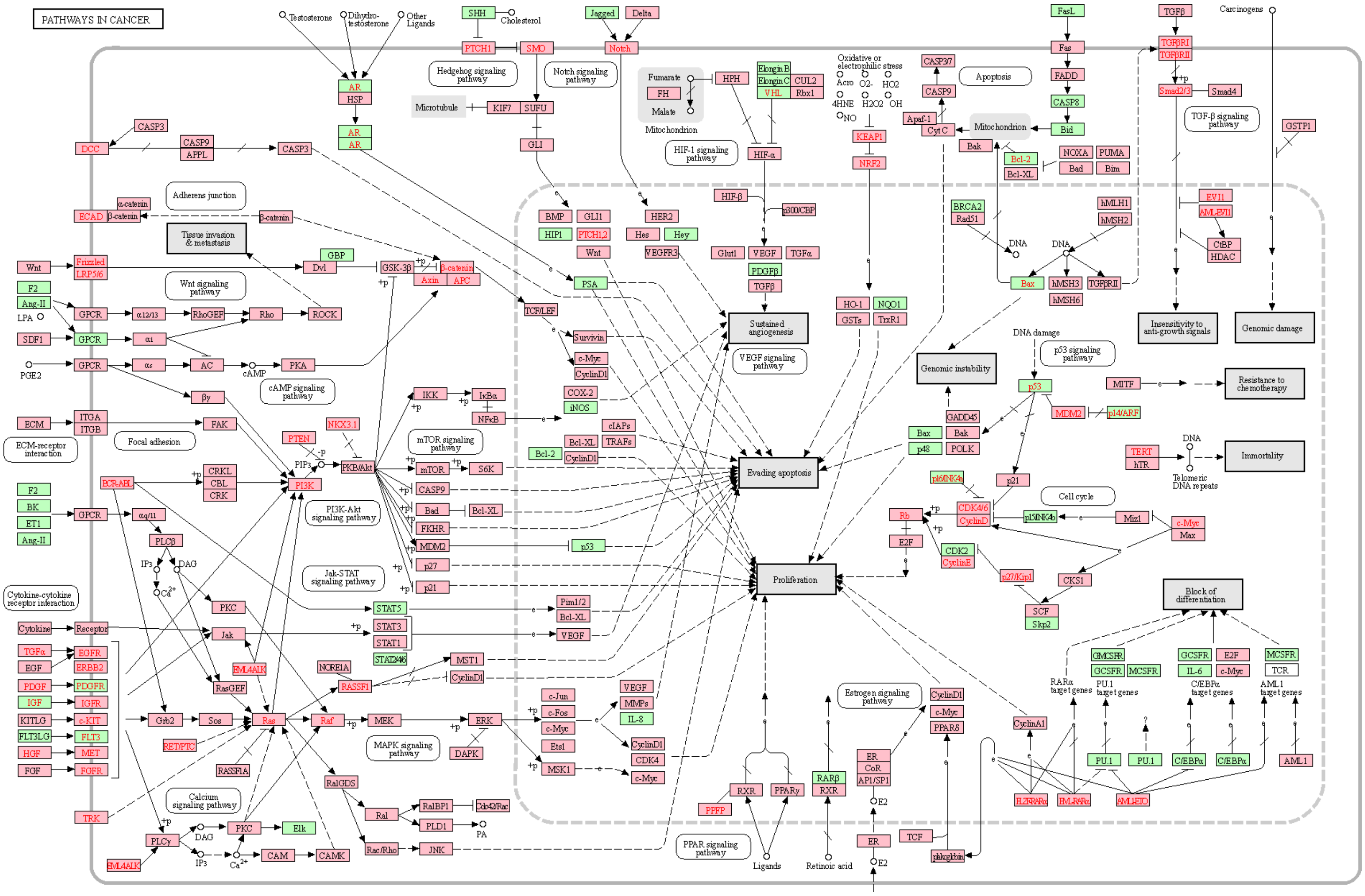


**Fig. S3e |** KEGG enrichment analysis of molecules involved in cancer pathways (pink) and represented in overlapping ChIP-Seq peaks in MT1 and MT2 clones normalized to WT cells.

**Supp. Fig. 4 |**

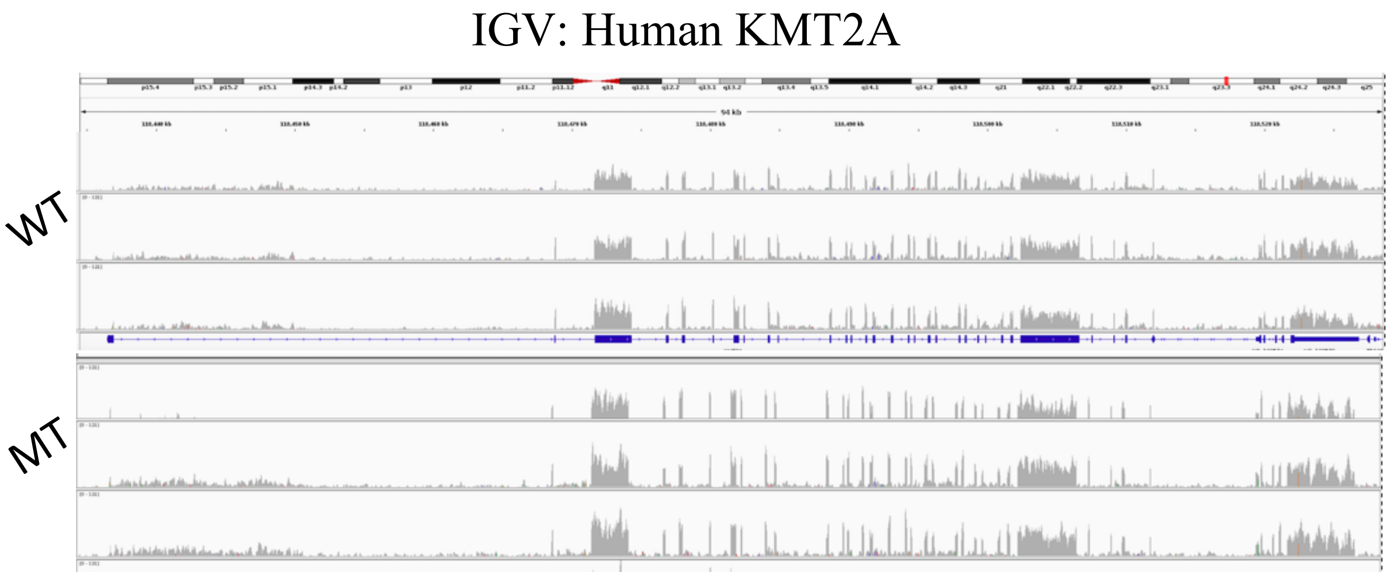


**Fig. S4a |** Integrated genomics view (IGV) of the human KMT2A locus in WT and MT (MT2) cells, showing identical transcript profile from multiple exons in triplicate RNA-seq samples.

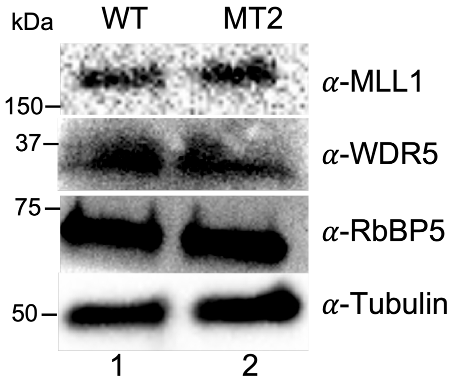


**Fig. S4b |** Protein levels of select MWRAD complex components in WT and MT2 cells. Whole cell extracts from and MT2 cells were immunoblotted using antibodies specific for MLL1, WDR5, and RbBP5. Tubulin was used as a loading control.

**Uncropped Western Images for Figures 2A, 5C, 6A and S3B.**

A cropped version of each figure is shown for reference. In some cases, a longer exposure is shown to help visualize the blot outline. Cropped areas of each blot are outlined in red.

Fig 2a uncropped images

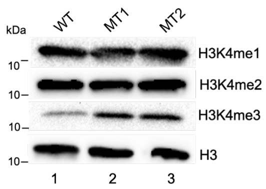


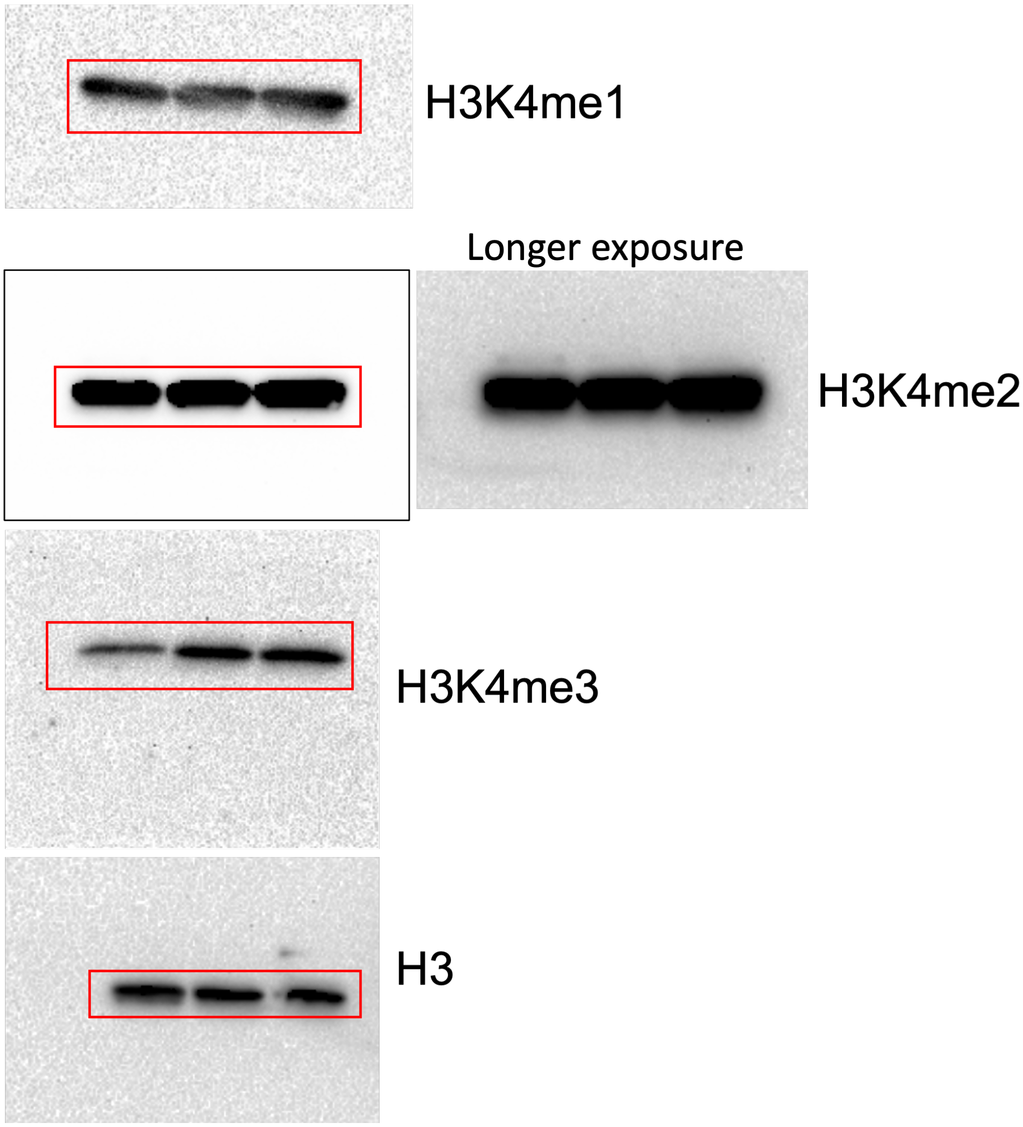

### Fig. 5C Uncropped Images

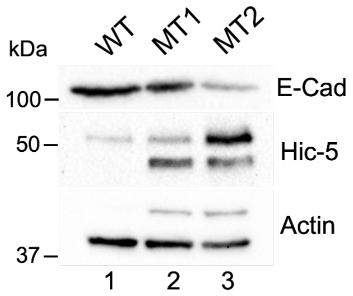


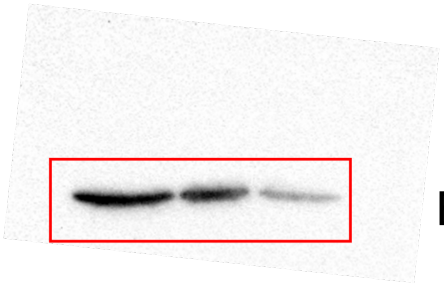

E-Cad

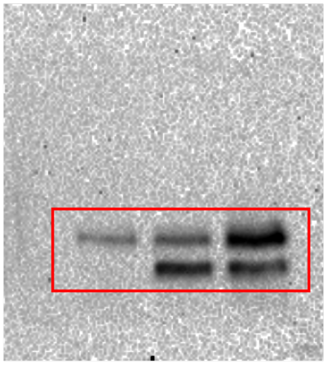

Hic-5

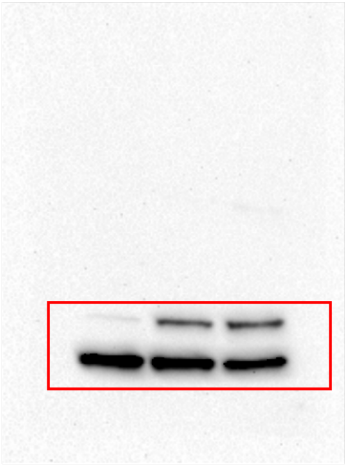

Actin

### Fig. 6A Uncropped Images

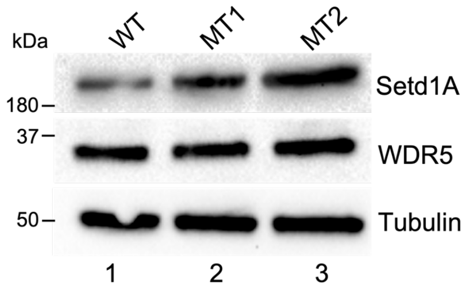


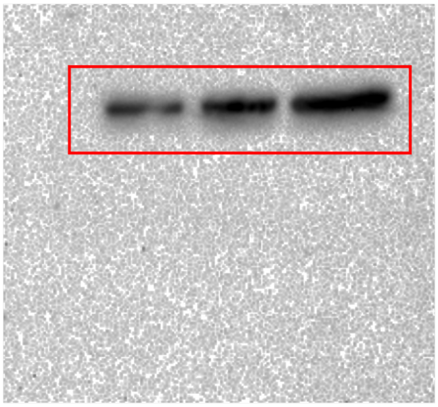

Setd1A

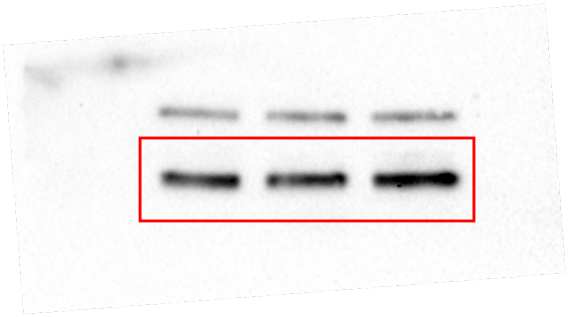

WDR5

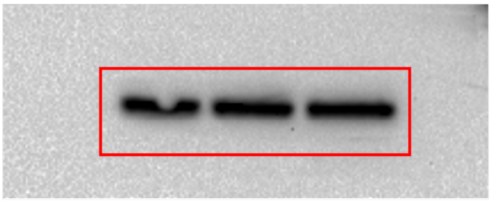

Tubulin

### Fig. S3B Uncropped Images

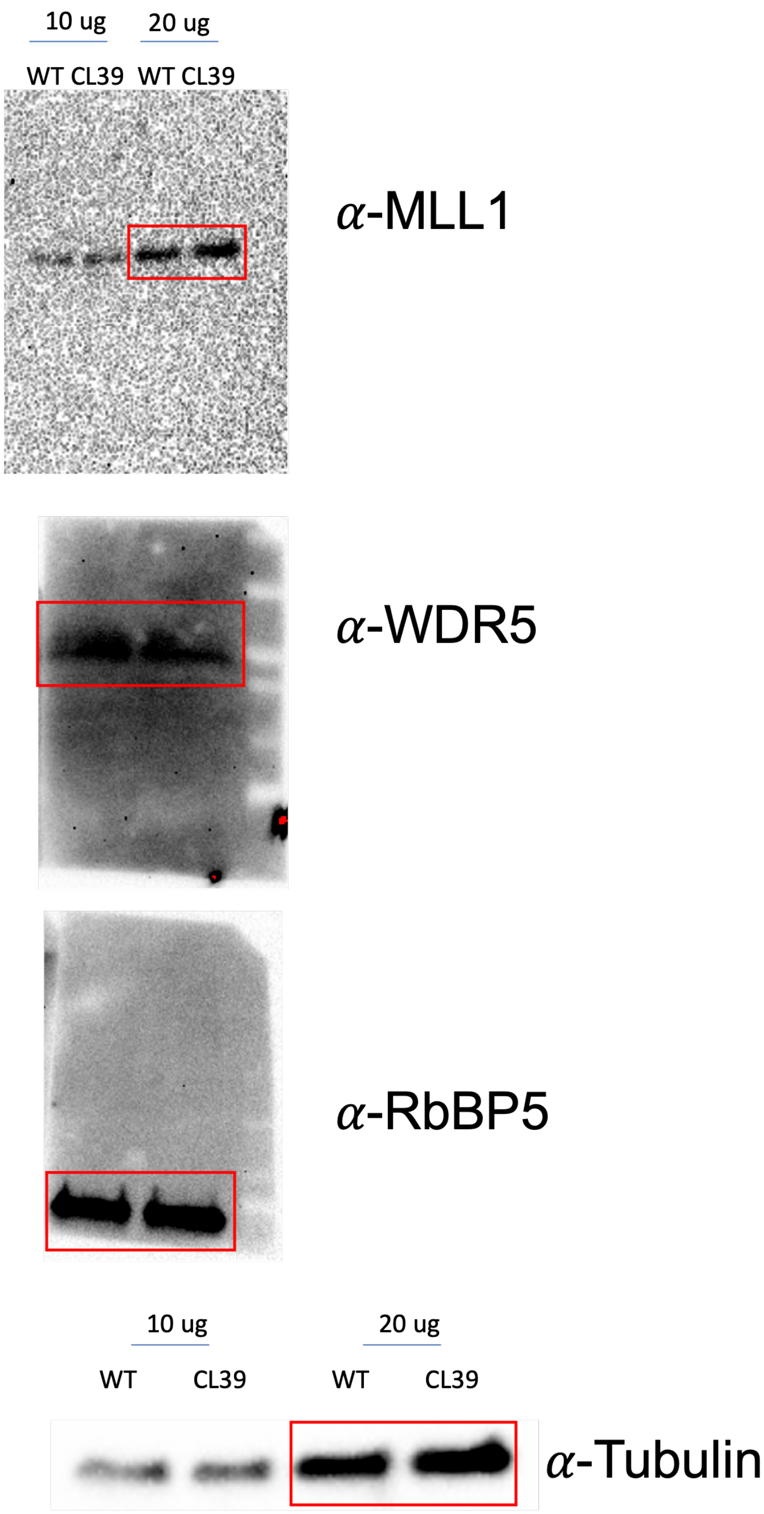


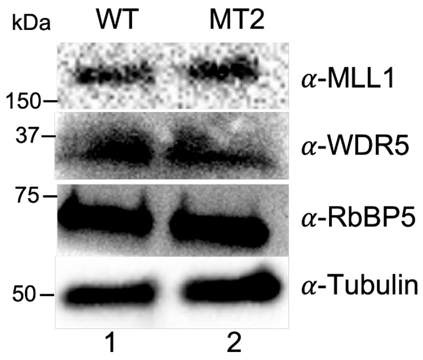

### Supporting Information

#### Antibodies and Primers

| **Primary Antibodies** | **Manufacturer, Catalog Number** | **Dilution Factor/Application** |
|---|---|---|
| Mouse anti-E-cadherin | BD Biosciences, 610182 | 1:1000 WB |
| Rabbit anti-E-cadherin | Cell Signaling, 3195 | 1:100 IF |
| Mouse anti-Hic-5 | BD Biosciences, 611165 | 1:1000 WB, 1:100 IF |
| Rhodamine-phalloidin (1:500, R415; Cat# PHDR1, Denver, CO) | Cytoskeleton, R415 PHDR1 | 1:500 IF |
| Rabbit anti-Histone H3 | Millipore, 06-755 | 1:2000 WB |
| Rabbit anti-H3K4me1 | Abcam, ab8895 | 1:1000 WB |
| Rabbit anti-H3K4me2 | Abcam, ab32356 | 1:5000 WB |
| Rabbit anti-H3K4 trimethylation | Active Motif, 39159 | 1:1000 WB |
| Rabbit anti-H3K4 trimethylation ChIP Grade | Epicypher, 13-0041 | 1 ug per ChIP |
| Rabbit anti-Setd1A ChIP Grade | Abcam, ab70378 | 1:2000 WB, 1 ug per ChIP sample |
| Rabbit anti- beta Actin | Abcam, ab8227 | 1:1000 WB |
| Mouse anti-Tubulin | Sigma, T5168 | 1:1000 WB |
| Rabbit anti-WDR5 | Abcam, ab22512 | 1:1000 WB |
| Mouse monoclonal anti-MLL1 | Millipore, 05-765 | 1:1000 WB |
| Rabbit anti-RbBP5 | Bethyl Labs, A300-109A | 1:1000 WB |
| Rabbit IgG negative control antibody for ChIP | Thermo Fisher Scientific, 31235 | 1 ug per ChIP sample |
| **Secondary Antibodies** | | |
| Goat anti-mouse DyLight 488-conjugated | Thermo Fisher, 35502 | 1:250 IF |
| Goat anti-mouse DyLight 550-conjugated | Thermo Fisher, 84540 | 1:250 IF |
| Goat anti-rabbit DyLight 488-conjugated | Thermo Fisher, 35552 | 1:250 IF |
| Goat anti-rabbit DyLight 550-conjugated | Thermo Fisher, 84541 | 1:250 IF |
| Secondary anti-rabbit IgG, HRP-conjugated | GE, NA934VS | 1:30,000 |
| Secondary anti-mouse IgG, HRP-conjugated | GE, NA931VS | 1:10,000 |

| **Primer Name** | **Forward Primer (5’>3’)** | **Reverse Primer (5’>3’)** |
|---|---|---|
| **ChIP-qPCR** | | |
| ZFP42 | ATTGGGCATTCCAGCCTACC | GAGTTCAGCTCCTTGGACCC |
| ACTIN | AGCCTCGCCTTTGCCGA | GCGCGGCGATATCATCATC |
| MMP1 | ATGCTGAAACCCTGAAGGTG | CTGCTTGACCCTCAGAGACC |
| MEIS1 | TGTGTAAGACGCGACCTGTT | CCGTGCGTGTGTAAAGTGTG |
| GAPDH | TACTAGCGGTTTTACGGGCG | TCGAACAGGAGGAGCAGAGAGCGA |
| HOXA9 | AGTGGTCTCACTTCCTCCCA | GCTGAGCAGTAAAAGGTGGC |
| **RT-qPCR** | | |
| DLK1 | GGGCACAGGAGCATTCATAG | GACGGGGAGCTCTGTGATAG |
| TBXT | GCCCTCTCCCTCCCCTCCACGCACAG | CGGCGCCGTTGCTCACAGACCACAGG |
| SOX17 | CGCTTTCATGGTGTGGGCTAAGGACG | TAGTTGGGGTGGTCCTGCATGTGCTG |
| GAPDH | ACATCGCTCAGACACCATG | TGTAGTTGAGGTCAATGAAGGG |
